## Supplemental Materials for "Pocket MUSE: an affordable, versatile and high performance fluorescence microscope using a smartphone"

#### Supplemental notes and figures

### Table of contents

**1. Basic design considerations** **1**
1.1 Reversed aspherical compound lens (RACL) 1
1.2 UVC LED 2
1.3 LED adapter 3
1.4 Sample holder (optical window) 4
1.5 D printed components 5
1.6 Assembly and alignment 7
1.7 Electronics 8
1.8 List of essential components 11
1.9 List of tools and supplies 12

**2. Advanced design considerations** **13**
2.1 Total internal reflection (TIR) coupling - geometric ray tracing 13
2.2 Total internal reflection (TIR) coupling - interface coupling efficiency 16
2.3 Angled coupling interface 17
2.4 LED triggering 19
2.5 Adjustable focusing 20
2.6 Smartphone/camera selection 21

**3. Using Pocket MUSE** **22**
3.1 Staining and dye selection 22
3.2 Sample loading and unloading 24
3.3 Data acquisition 25
3.4 Raw data conversion 26

**4. Additional characterization and images** **30**

**5. References** **40**

### 1. Basic design considerations

Pocket MUSE includes many non-standard parts from various e-commerce vendors. There are dimensional variations between parts acquired from different sources. Availability of these components also varies in different regions of the world. In addition, as some of the original components require modifications, several details of the actual mechanical design needs to be adjusted accordingly. Therefore, instead of showing a fixed blueprint of Pocket MUSE, we aim to provide detailed guidelines for designing and building a Pocket MUSE from scratch. Following instructions provided in this section, users with some basic prototyping knowledge should be able to identify, modify, design and make each sub-component of Pocket MUSE, and assemble them into a functional device.

##### 1.1 Reversed aspherical compound lens (RACL)

RACL [1] is one of the most critical components in Pocket MUSE. During our early stage prototyping, we were able to acquire some lens samples from an OEM lens manufacturer (Largan Precision). However, possibly due to the recent high demand of smartphone cameras, the company regularly ignored inquiries for small sample orders recently. Therefore, we concluded that the most reliable way to obtain a small amount of RACLs for Pocket MUSE is from aftermarket or replacement camera parts for mobile devices, especially from camera modules using 1/6.5” - 1/8” image sensors (e.g., a laptop webcam).

Based on our preliminary research, most mainstream laptop webcams after 2016 share similar technical specifications. With f-number <2.2 and 4-element optical designs, most of these lenses can provide optical resolution down to 1 μm and well-corrected aberrations over 2/3 FOV. As the field angles of most webcam lenses are around 70-75°, when used as RACL objectives, magnifications of the lenses are proportional to sizes of the camera sensors.

It is difficult to simulate the exact optical design of the lenses because a lot of the information is not disclosed to the public by the manufacturers. Therefore, as long as desired imaging performance is achieved by testing the lens using a resolution target, it is unnecessary to determine the exact number of these specifications for a DIY Pocket MUSE. For reference, we have verified that the webcams for various Lenovo laptops (e.g., Part# 04X1399, 04X1398, 04X0294, 04X0273 and 04X0297) works with Pocket MUSE. We have not verified, but based on preliminary research, some Dell (e.g., Part# Y2TKG) and HP (e.g., Part# 833474-1K0) laptop webcams should also work. Cost of these replacement cameras varies from $3-20, and they could also be found in used/broken laptop computers.

In the original packaging (**Supplemental Figure (SF) 1.1**), the lenses are usually installed in the webcam housing with screw based fasteners. To detach a lens from the device and avoid mechanical deformation, it is recommended to gently unscrew it using a pair of plastic tweezers. Since the optical elements are molded with plastic, they are very susceptible to scratching damage. To prevent scratching the lens surface, it is recommended to cover the front surface of the lens with a piece of tape during any handling. Still, some accidental minor scratches will not significantly affect the image quality for most tasks, but it should be avoided to ensure the best performance.

---

**Supplemental Figure 1.1**
RACL detached from a 04X1399 Lenovo laptops

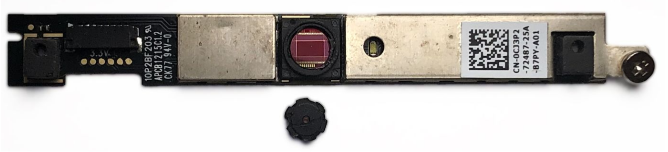

#### 1.2 UVC LED

LEDs at the UVC wavelength range [2] are becoming increasingly available in recent years. They are primarily used in various disinfection and sterilization applications. At the time, these LEDs can be found in various independent vendors on eBay in small quantities and relatively low cost. To search for an appropriate UVC LED for Pocket MUSE, a good keyword could be "275 nm LED" or "285 nm LED".

UVC LEDs often have significantly different packages [3] from other typical visible and near UV LEDs. The majority of the UVC LEDs are made of inorganic materials, because most plastic materials are highly absorptive at the UVC wavelength range. For Pocket MUSE, the most suitable (smallest) UVC LED package is often referred to as "SMD3535" or simply "3535", meaning the XY dimension of the LED package is 3.5x3.5 $mm^2$. One such LED package often comprises an AlGaN UVC LED die/chip (a flip-chip surface mount device (SMD), **SF 1.2**), a transient-voltage-suppression (TVS) diode, an AlN-PCB based housing with 3 SMD pads (cathode/heatsink/anode), and a quartz (or fused silica) protection window.

For Pocket MUSE, the LED die needs to be placed right next to the side of the sample holder (optical window). Therefore, it is necessary to remove the protection window and reduce the height of the dam (side enclosure) of the original LED package. The protection window could be removed from the original UVC LED package by prying using a utility knife. After that, the dam of the housing should be polished down to the level of the LED die using a ~200 grit file or sandpaper. Example photo of the LED, cross-section structure and the modification is shown in below.

In addition to wavelength and package size, other key technical specifications of the LEDs include optical power (2-5 mW), voltage (6-7.5 V) and current (30-150 mA). A brighter/higher power LED often provides shorter exposure time and better signal to noise ratio, but it can also lead to shorter battery life, more heating (shorter LED life) and higher cost. Thus, with high performance smartphone cameras, lower power LEDs are prefered. Cost of these UVC LEDs varies from $2-15 on e-commerce, and is continuously dropping.

---

**Supplemental Figure 1.2**
Chip structure [4] (top left), package (top right) and modification (bottom) of a typical UVC LED for Pocket MUSE

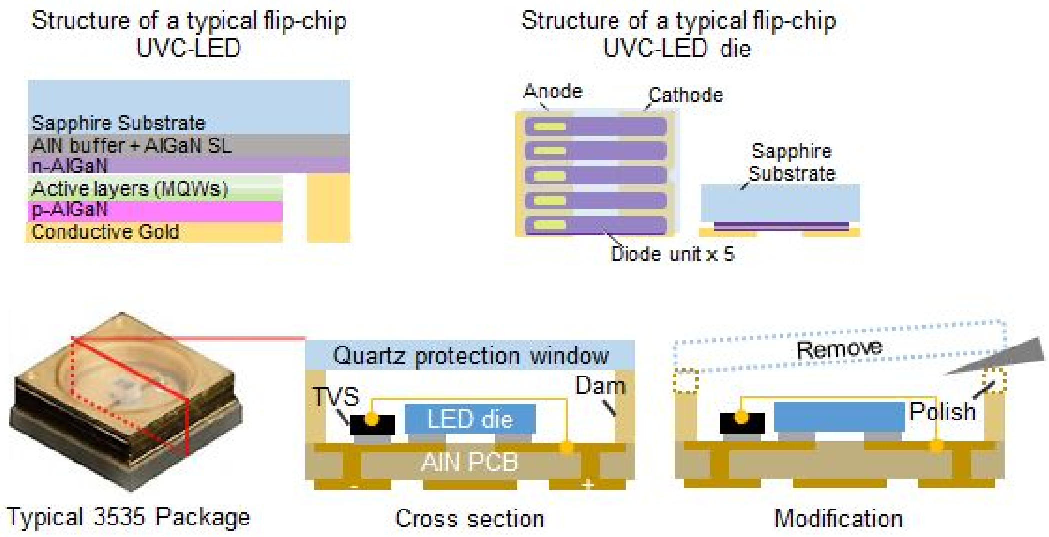

#### 1.3 LED adapter

Although it is feasible to power the LEDs directly with electric wires, it is highly recommended to mount the LED on a piece of PCB for better heat dissipation and ease of mounting. As shown below, the primary function of the PCB is an adapter for the LED and electric wires. The two pads near the edges of the PCB are connected to the cathode and the anode of the LED. Two thru-holes (for electric wire soldering) are located at one end of the PCB. A screw mounting hole is located at one end of the middle mounting/heatsink pad. The LED can be soldered on the PCB using a laboratory heat plate or a low cost benchtop reflow oven. The assembled PCBs are inserted in the baseplate by press fitting, and a metal screw that further secures the attachment. The screw also serves as a heart sink to provide extra heat capacity.

The LED adapter PCB is also easy to fabricate. Because the design only contains holes and straight lines, it is simple enough to be fabricated manually from blank FR4 copper clad PCB using a ruler, a marker (draw the cuts), a utility knife (cut the board and scrap the gaps between the trace (e.g., green lines in **SF 1.3**), and a power drill (drill the 3 holes). A thicker copper layer (e.g., ≥2 oz) is desired for better heat dissipation. Alternatively, the PCB LED adapter can be easily drawn with common PCB design software such as Autodesk Eagle, and manufactured with a benchtop CNC router or online PCB prototype services such as OSHPark and PCBway, but it may lead to higher cost and unnecessary large production volume (e.g., $15/50 boards). This option is more beneficial when combined with other customized  PCB (e.g., current supply).

---

**Supplemental Figure 1.3**
Design considerations for the LED adapter PBC, showing conceptual designs (top) CAD drawing (bottom left), and a fabricated assembly (bottom right)

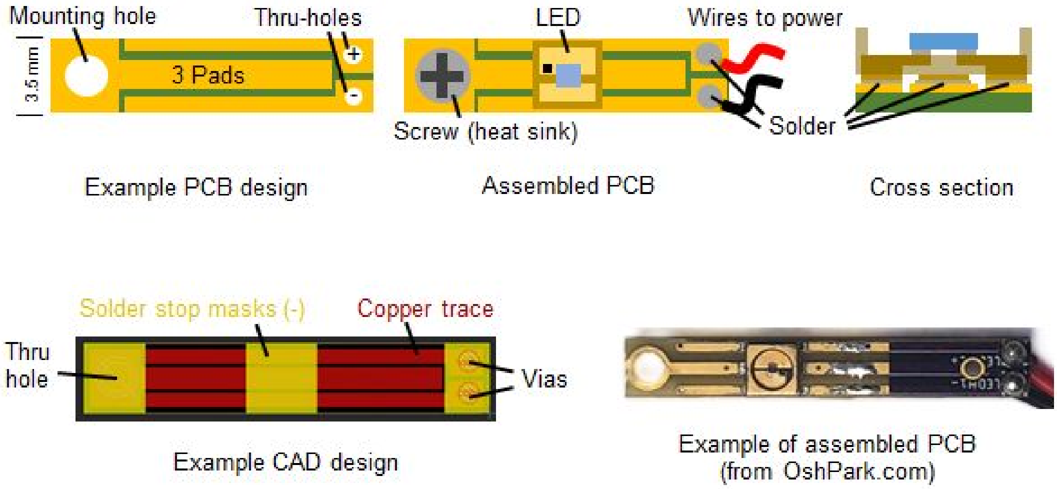

#### 1.4 Sample holder (optical window)

Most common optical materials such as borosilicate glass (e.g., BK7) are not transparent in the UVC range. Only a few common optical materials, including quartz, silica, sapphire and some UVC transparent plastics [5], can be used for MUSE imaging. Among these, sapphire is widely used in benchtop MUSE systems, but it is difficult to process sapphire without special tools and its high refractive index can lead to TIR illumination complications. UVC transparent plastics (e.g., polyolefins) are cheaper and easy to process, but they often have poor durability and chemical resistance. Therefore, sapphire and plastic optical windows should be avoided for the Pocket MUSE sample holder.

The other two materials, quartz (e.g., G1) and silica (e.g., S1-UV), are both good options for the sample holder. However, these materials may still have different optical grades, and some fraudulent vendors may falsely mark borosilicate as silica, especially for the products at the lower cost range. Therefore, it is necessary to verify the UVC transparency of the optical window. A simple quick test could be performed by illuminating a piece of white printing paper with a UVC LED in the dark. Brightness of the fluorescence from the paper should be minimally affected when the optical window is placed in between the paper and the LED.

0.5±0.1 mm thick quartz and silica optical windows are ideal starting points for the sample holder. They can be purchased from various scientific glass equipment vendors, including Ted Pella, Esco Optics, etc, usually at $10-20 per piece (usually in good quality). Alternatively, they could also be found on independent vendors on e-commerce (with lower price and more diverse quality).

Off-the-shelf optical windows are often too large (≥19x19 mm$^2$) for the Pocket MUSE sample holder (<10x10 mm$^2$). Therefore, one piece of optical window is divided into multiple Pocket MUSE sample holders as shown in **SF 1.4** below. To process the sample holder, first the optical window is manually cut with either a diamond scribe or a Dremel sanding disk. Size for the sample holder can be flexible. For instance, it is convenient to cut a 19x19 mm$^2$ optical window into 4 ~9x9 mm$^2$ smaller pieces, and 25.4x25.4 mm$^2$ into 9 ~8x8 mm$^2$ pieces. After cutting, the optical windows should be hand polished into more accurate sizes using 300-1000 grit sandpaper. To ensure the most efficient LED coupling, two opposite sides (e.g., 0.5x9 mm$^2$ faces) of the optical window are further fine polished with 40/30/12/9/3/1/0.3 μm lapping films. To prevent scratches on the optical window, it is necessary to protect the glass surface with tape during the process.

---

**Supplemental Figure 1.4**
Modification of an off-the-shelf optical window into a Pocket MUSE sample holder

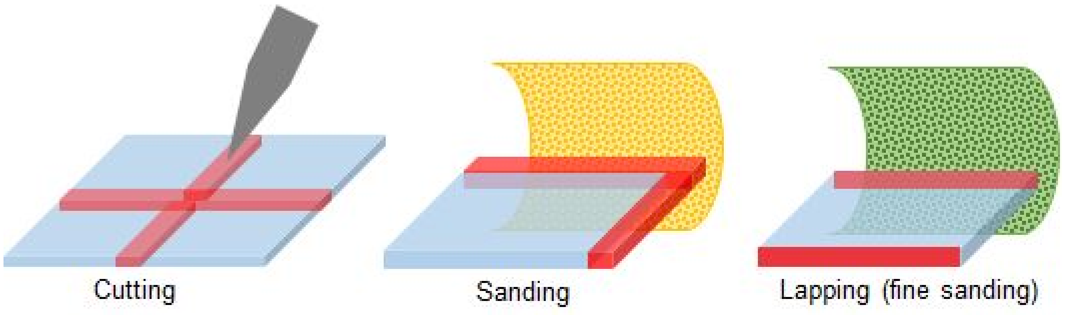

#### 1.5 3D printed components

Pocket MUSE has 2 major 3D printed mechanical components (**SF 1.5**) : a baseplate and a sample holder retainer, both can be printed with simple fused deposition modeling (FDM) 3D printing [6]. Their primary function is to provide the integration mechanism for the rest of the components. The mechanical design concept of Pocket MUSE is shown below. Because some fine features (e.g., hole diameter, gap width, etc.) require high dimensional accuracy (e.g., ±0.05 mm), it is necessary to measure the dimensions of each component (e.g, thickness of the sample holder) before designing the 3D model. In most cases, it is also necessary to fine-tune the small dimensional features with a few 3D printed samples before finalizing the design.

Although consumer grade FDM printers often have poor initial accuracy, precision (e.g., ±0.05 mm) and repeatability are usually sufficient. Therefore, by measuring the dimensional errors in the sample, it is possible to offset these errors in the modified designs. In addition, most 3D printers have limited layer height options due to discretized steps of linear stages. Thus, the feature sizes in the Z-dimension should be an integer multiple of the step size. Printing at a thinner layer thickness is unnecessary because focus alignment of Pocket MUSE does not rely on 3D printing accuracy in the vertical dimension. Based on our experience with a hobbyist FDM 3D printer (e.g., Snapmaker), the most reliable layer thickness is 0.1 mm, but this will vary for different 3D printers.

Because the demonstrated version of Pocket MUSE was designed for low-volume production with FDM 3D printing, complex fine features are avoided in the design. Instead, screw-based fastening mechanisms are used to secure the sample holder retainer. In case high-resolution 3D printers are available (e.g., stereolithography, jetting, etc.), screw-based fastenings could be replaced by snap fit designs for more convenient assembly. Still, the screws are recommended for the LED PCBs for thermal management purposes.

---

**Supplemental Figure 1.5**
Mechanical design concept of Pocket MUSE.

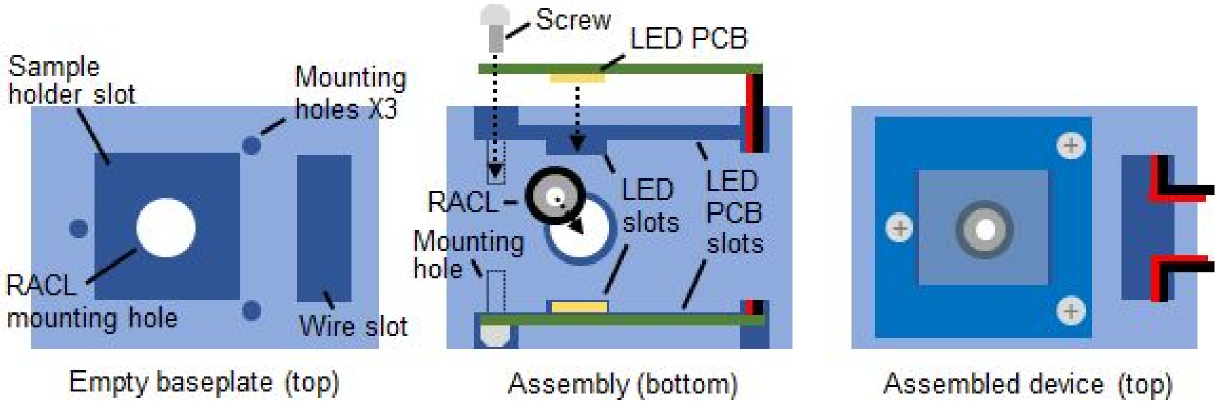

An example design of the baseplate and the retainer is shown in the diagrams (**SF 1.6**) below. The body of the baseplate is a ~20x35x5 $mm^3$ rectangular block. The top face of the block includes a sample holder slot, a wire slot and optional mounting holes for the sample holder retainer. A thru-hole for RACL mounting is located at the center of the sample holder slot. The bottom face of the block includes two LED PCB slots. The physical dimensions of these features need to meet the following design rules : **1.** The XY dimensions of the sample holder slot should allow a loose fit of the sample holder window and the retainer; **2.** The depth of the sample holder slot should be set, so the back focal plane of the RACL is ~0.1-0.2 mm below the sample holder surface; **3.** The thru-hole geometry should be set, so the RACL can be held with friction; **4.** The LED slots should be aligned to the edge of the sample holder slot, so the slots are connected at the interfaces; **5.** The depth of the LED PCB slots should be set, so the LED chips are approximately at the center lateral plane of the sample holder window; **6.** The widths of the LED PCB slots should be set to allow frictional fit of the LED PCBs; **7.** The centers of the LED chips should be approximately at the transverse center line of the sample holder; **8.** The wire slot and the LED PCB slots should be connected, and two chamfers should be created at the bottom of the wire slot (on edges next to the LED PCB slots); **9.** The height of the retainer is set, so the sample holder window is held against the bottom of the sample holder slot; **10.** The screw mounting holes should be tapped for small screws (M2x0.4 (metric) or 2-56 (imperial)). Location of the feature rules are labeled in the drawings below.

---

**Supplemental Figure 1.6**
Example 3D drawings of the 3D printed parts of Pocket MUSE

.

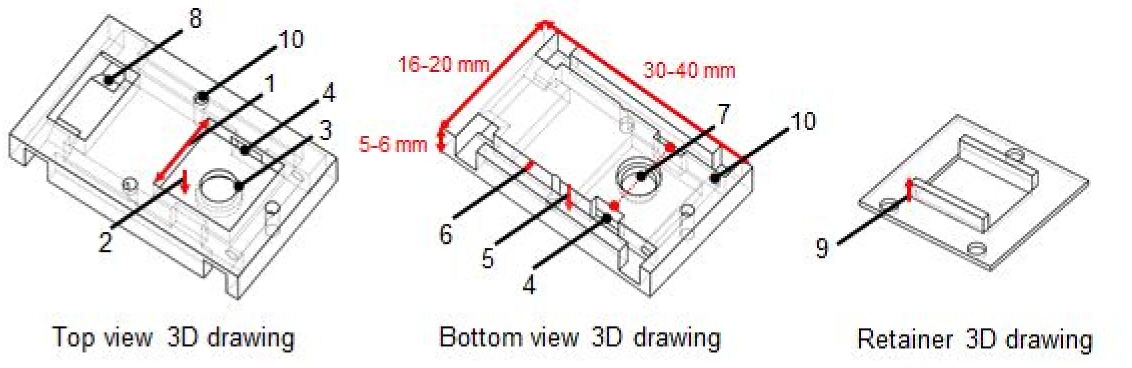

#### 1.6 Assembly and alignment

The major assembly steps are shown in the diagram below. In brief, the LED PCBs are inserted into the bottom LED PCB slots by press fitting. A screw may be used to secure the attachment as shown in **SF 1.3 & SF 1.5**. The RACL is inserted into the RACL mounting hole by press fitting. This is achieved by loosely inserting the RACL in the mounting hole, and pushing it in against a flat surface. The sample holder window is inserted in the sample holder slot and held by the retainer (**SF 1.7**).

The iterative alignment process of Pocket MUSE is shown in the diagram (**SF 1.7**) below. To start, the initial assembled device is attached to a smartphone. An out-of-focus image of debris and dust particles on the surface of the sample holder can be visualized on the camera preview. This should provide a qualitative assessment of the amount the sample holder surface is out of focus. Next, the sample holder and the RACL (gently pushed out with plastic tweezers) are removed from the base-plate. After checking the thickness of the sample holder using a caliper, the bottom of the sample holder is thinned with 1000-3000 grit sandpaper. Use the caliper to measure the thickness regularly to prevent over sanding. After removing ~30-50 µm of material, the device should be assembled and attached back to the smartphone to verify if the image is in focus. The initial layer thickness of FDM printing can vary by ±50 µm. For a 3D design in which the RACL focal plane is ~100-200 µm below the sample holder surface, it is necessary to repeat the iterative process 3-5 times. A sharp image of the sample holder surface is an indication of good alignment. High accuracy (e.g., ±20 µm) of the thickness is not required because the smartphone camera autofocus. When aligning for the first time, one might overdue the sanding, but should get a good idea of what the optimal thickness and image sharpness will be for future alignments.

---

**Supplemental Figure 1.7**
Assembly (top) and alignment (bottom) of Pocket MUSE

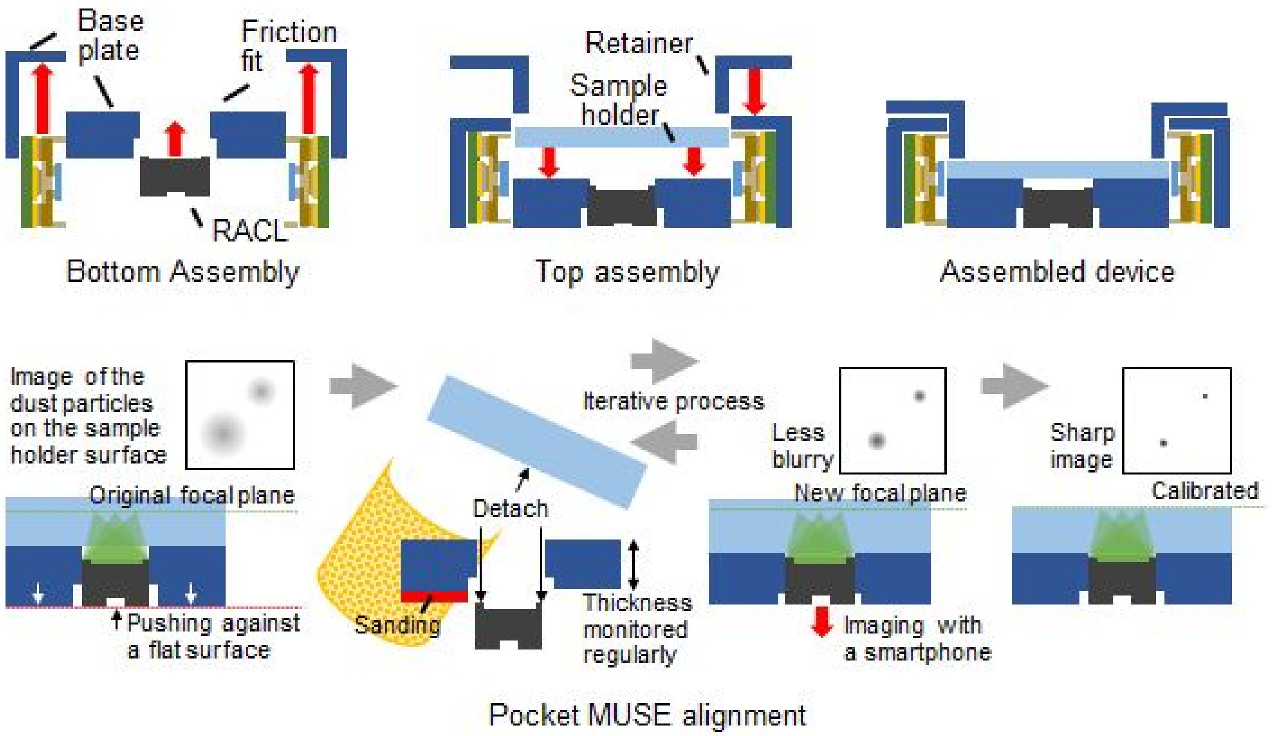

#### 1.7 Electronics

The primary function of the Pocket MUSE electronics is to supply power and provide an enabling mechanism for the UVC LEDs. There are multiple ways to drive the UVC LEDs in Pocket MUSE. For each method, there are trade offs between cost, physical size, reliability, battery life and prototyping complexity. It is often recommended to drive a UVC LED at the rated current. However, when long term reliability is not a concern, the LEDs could also be driven with constant voltage directly. Selection of the power supply method should be also determined based on the actual user needs and the level of prototyping skills.

UVC LEDs often have rated voltages around 6.5-8 V. For either current driving or voltage driving, it is necessary to start with a power supply above the rated voltage. The easiest way to drive Pocket MUSE is to use a few primary cells (non rechargeable batteries) in series or a 9V battery directly. For instance, 4-5 AAA batteries (1.5-1.65 V) connected in series can be used to drive the LEDs directly. In case the voltage is above the rated voltage of the LEDs, a simple way to reduce the supply voltage is to connect a few diodes (e.g., 1n4001) in series, and each diode can drop the voltage by 0.7 V.

Despite the simplicity of the design, there are several disadvantages in using the primary cells directly. Since primary batteries are not rechargeable, they are not eco-friendly and can cost more in the long run. Alternatively, it is possible to connect two 3.7 V lithium-ion rechargeable batteries in series to achieve the 7.4 V supply voltage. However, in either case, the footprint of multiple batteries is many times bigger than the Pocket MUSE itself. Voltage drops over time also causes gradual dimming of the illumination.

A slightly more complicated approach is to use a single lithium-ion rechargeable battery with a DC-DC up-regulator (boost converter), which converts 3.7 V to a desired higher DC voltage (**SF 1.8**). This approach provides several benefits over the direct battery driving strategies, including small footprints, lower cost and stable voltage supply. DC-DC up-regulator boards could be found at most electronic vendors (e.g., Adafruit) at <$5. Commonly-found 6 V, 9 V or variable DC-DC up-regulators are all appropriate options for Pocket MUSE. Typical UVC LEDs usually have rated voltages at 6.5-8 V, but 6 V may provide sufficient optical power (e.g., >1 mW) for Pocket MUSE. To achieve the highest rated optical power (e.g., >3 mW), it is possible to use a variable DC-DC up-regulator (large footprint), or use a 9 V regulator and drop the voltage to the desired value with diodes (higher power consumption).

---

**Supplemental Figure 1.8**
Constant voltage driver circuitry for Pocket MUSE.

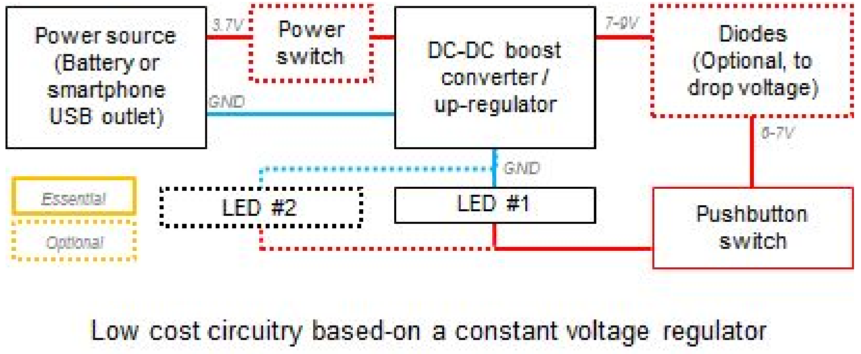

For the highest reliability, it is still recommended to drive each UVC LED at the rated current using a constant current supply (**SF 1.9**). However, because compact, constant current supply circuits are not readily available for order from electronic vendors, it is necessary to customize the circuit from scratch. Simple current drivers could be built around LM317 regulators, but some entry level PCB design skills are required. Since LEDs are connected in parallel (to prevent doubling the supply voltage), each driver should contain two independent current sources. These current sources could be supplied with a 9 V battery or a DC-DC regulator. For production, current driver PCBs could be panelized with the LED adapters.

For some smartphones with sufficient battery capacity (e.g., >2500 mAh), it is also possible to outsource the power from the smartphone battery directly through the USB or Lightning port (**SF 1.10**). For micro-USB ports, power can be outsourced from the ports directly with a micro-USB male connector. For USB-C and Lightning ports, special OTG boards/chips are required to outsource from the device. One common source for OTG outsource connectors is from smartphone-powered mini portable fans. Alternatively, they could also be purchased from OEM vendors at large volumes (e.g., >100 pieces). Using the smartphone battery essentially eliminates the need for an external battery and charging equipment, which in turn reduces the cost and the footprint of the system. However, it also sacrifices the battery life and health over time, especially when a large current is sourced (e.g., >150 mA) with high power LEDs and low efficiency drivers. Therefore, this approach should only be considered after carefully balancing the user needs and disadvantages.

For each of these power supply schemes, a pushbutton switch is connected in between the LEDs and power supply, which serves as the enabling mechanism. In addition to the pushbutton switch, we also recommend installing a power switch before the DC-DC up-regulator to avoid power loss during standby and unprecedented enabling of the device.

---

**Supplemental Figure 1.9**
Constant current driver circuitry for Pocket MUSE.

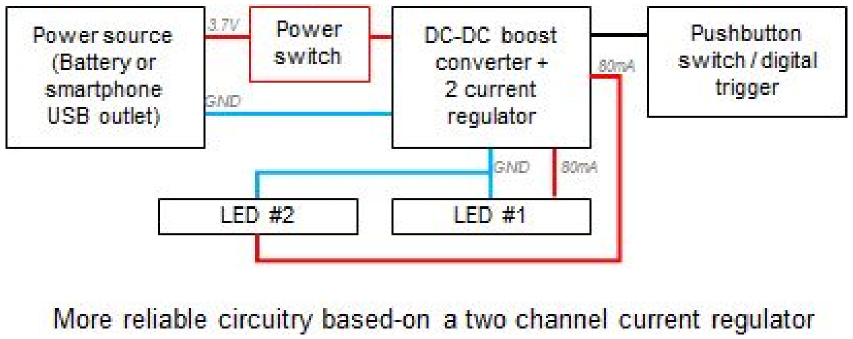

**Supplemental Figure 1.10**

Additional images showing the circuitry designs of Pocket MUSE

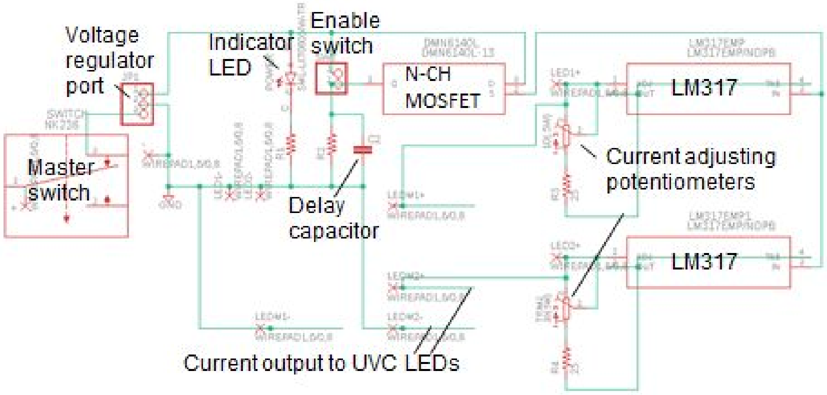


Example circuit design of a 2-channel current regulator

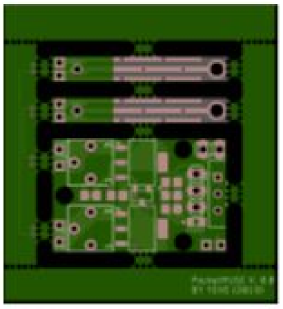

Example PCB panel CAD of a customized current regulator with 2 LED adaptors

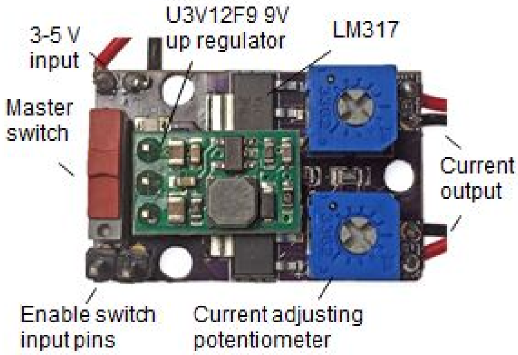


Example assembled 2-channel current regulator

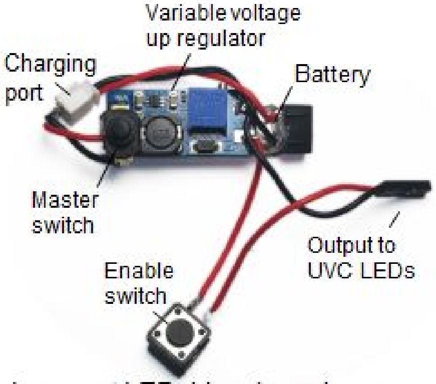


Low cost LED driver based-on a constant voltage regulator and an external battery

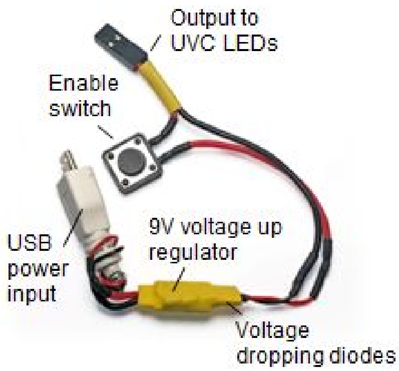


Low cost LED driver based-on a constant voltage regulator and USB power input

#### 1.8 List of essential components

**Supplemental Table 1.1**
List of essential Pocket MUSE components (for each Pocket MUSE device)

| Component | Cost ($) | Quantity | Source / supplier | Remarks |
|---|---|---|---|---|
| Reversed aspherical compound lens (RACL) | 3 - 20 | 1 ea | eBay / Alibaba / local electronics (flea/used) market / old laptops | Detached from laptop webcams with 1/6.5” - 1/8” sensors |
| 265-285 nm UVC LED | 1 - 15 | 1 - 2 ea | eBay / Alibaba / DigiKey / Mouser | Protection window removed / side dam polished to the die level |
| LED adaptor (PCB) | 0.1 - 5 | 1 - 2 ea | Blank FR4 copper clad PCB / OshPark / PCBway | Band-made or customized from PCB fabrication services |
| Sample holder (quartz / silica optical window) | 2.5 - 15 | 1 ea | Ted Pella / Esco Optics / Alibaba | Cut into smaller pieces |
| Base plate and retainer | <0.1 | 1 ea | 3D printed | PLA (FDM) or resin (SLA / jetting) |
| Wire | <0.1 | 1 ft | Any electronic supplier | < 26 AWG |
| Batteries and casing | 0.2-5 | 1 ea | Any electronic supplier | >3.3 V / >200 mA |
| DC-DC boost converter / up regulator (optional) | 3-6 | 1 ea | Pololu / Adafruit / eBay / Alibaba | Variable output or fixed 6 - 9 V output |
| Diode (optional) | <0.1 | 0 - 3 ea | Any electronic supplier | Voltage drop 0.7 V |
| Pushbutton Switch | <1 | 1 ea | Any electronic supplier | |
| Constant current driver (optional) | 10 - 50 | 1 ea | Customized | Not recommended for low volume production |
| Smartphone OTG power outlet (optional) | 5-10 | 1 ea | eBay / Amazon / Alibaba | From smartphone-powered mini portable fans |
| M1.6 / M2 / #2-56 screws | < 0.5 | 2 - 5 ea | Any hardware supplier | |
| Double sided tape | < 0.1 | 1X0.5 $in^2$ | Any hardware supplier | Rubber-based adhesive prefered |

#### 1.9 List of essential tools and supplies

**Supplemental Table 1.2**
List of essential tools and supplies for fabricating Pocket MUSE. Make sure you have access to the listed tools and supplies before get started

| Tool | Remarks |
|---|---|
| Tweezers | Handling fine components |
| Utility knife | Cutting PCB / modify 3D printed parts |
| Permanent marker (fine) | Cutting PCB |
| Ruler | PCB fabrication |
| Caliper | Fine measurements / focus calibration (thickness) |
| 3D printer | Baseplate & retainer fabrication / FDM / SLA / jetting / online service optional |
| Soldering tools | Soldering iron / magnifying glass / helping hands |
| Lab hot plate / reflow oven | Soldering surface mount devices (LED to led adaptor) |
| Common electronic supplies | Solder / solder paste / flux / Dupont connector / heat shrink tubing / electrical tape |
| Tap tool | Tapping screw mounting holes / corresponding to the screws |
| Office tape / masking tape | Glass surface protection |
| Sandpaper | Focus calibration / LED modification / 200, 300, 1000, 1500 & 3000 grit |
| Diamond scribe | Cutting glass |
| Lapping films | Fine polishing (sample holder edge) 40/30/12/9/3/1/0.3 μm |
| Dremel tool / power drills | PCB fabrication / sample holder glass fabrication |
| Hex key / screwdriver | Retainer & PCB attachment |

#### 2. Advanced design considerations

Following instructions provided in the previous section, users should be able to fabricate a basic Pocket MUSE system with most functionalities. In this section, we further discuss some design considerations for more advanced Pocket MUSE systems. This additional information should help users gain a better understanding of the underlying design principles of Pocket MUSE, and allow users to improve the basic Pocket MUSE design to achieve better performance and more advanced functionalities. It is worth mentioning here that some advanced design considerations discussed in this section are nonessential and would lead to greater design complexity, fabrication difficulty, and higher cost. Therefore, these instructions are only recommended for projects that require cutting edge performance (e.g., high illumination efficiency, long battery life and higher resolution).

##### 2.1 Total internal reflection (TIR) coupling - geometric ray tracing

Effectiveness of Pocket MUSE illumination is influenced by multiple factors. In addition to the optical power of UVC LEDs, the amount of light delivered to the sample is also directly related to the coupling efficiency of the light into the sample holder (optical window), the amount of light that undergoes total internal reflection at the air-glass interface, and the amount of light that refracts out at the sample-glass interface. Here, we only discuss a few major factors that significantly affect coupling efficiency of the UVC LED into the sample holder.

Refraction and TIR [7] are two important factors that affect LED coupling and illumination efficiency of Pocket MUSE. Based on Snell's law, some basic geometric ray tracing (**SF 2.1**) can help to explain this process better. Assuming the LED is a point source in air and the sample holder is a rectangular prism, the light will propagate within the parallel glass interface by TIR at a small incidence angle. The minimal incidence angle to enable TIR is determined by the refractive indices of the rectangular prism and the material on top of the prism

---

**Supplemental Figure 2.1**
Derivation of the acceptance angle based on Snell's law and schematic showing conditions of frustrated TIR, critical angle and TIR.

$$\frac{\sin(\theta_1)}{n_2} = \frac{\sin(\theta_2)}{n_1}, \qquad \frac{\sin(\theta_3)}{n_2} = \frac{\cos(\theta_2)}{n_3},$$

$$\frac{n_1^2\sin^2(\theta_1) + n_3^2\sin^2(\theta_3)}{n_2^2} = \sin^2(\theta_2) + \cos^2(\theta_2) = 1,$$

$$n_1 = 1, \qquad \theta_3 = \frac{\pi}{2}, \qquad \sin(\theta_{1/TIR}) = \sqrt{n_2^2 - n_3^2}$$

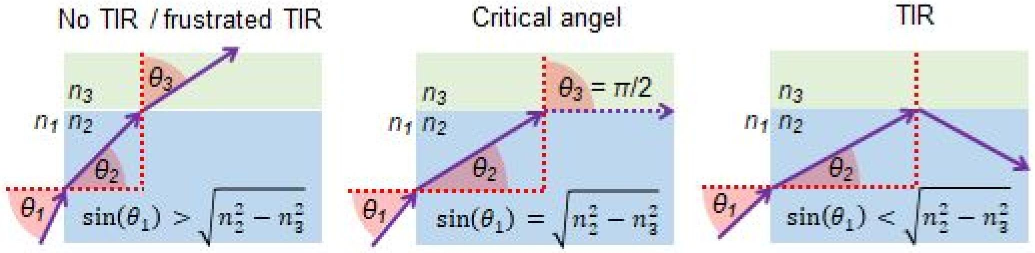

The overall goal of the frustrated TIR design is to ensure the majority of the light is coupled into the glass, propagates through the glass by TIR, and frustrates out after reaching the sample. As shown in the diagram (**SF 2.2**) below, at the glass-air interface, total internal reflection would happen when the coupling angle is below a certain value, which is determined by the refractive index of the glass substrate. Because $\sin(\theta_1)$ is always smaller than 1, when the refractive index of the substrate is larger than 1.414, light coupled into the glass substrate from the side face will always undergo TIR at the glass-air interface. Since most common glass substrates (e.g., quartz and fused silica) have refractive indices significantly greater than 1.414, TIR at the glass-air interface will always take place as long as the light is coupled into the glass from the side face. In other words, the acceptance angle is 0.5π for any glass materials. In reality, TIR is always a little lossy because the glass surface is not perfectly flat causing some light to scatter out of the glass. Some UV light would also be absorbed by impurities in the glass substrate. Therefore, it is beneficial to use a relatively small glass window to minimize power loss during the TIR process.

After the light reaches the sample-glass interface, there are potentially three ways it interacts with the sample. First, roughness of the glass at the sample-glass interface would cause a small amount of light to scatter out of the glass. Second, similar to frustrated TIR sensing, some light would refract out of the glass due to the change of the refractive index. Third, similar to TIRF microscopy, evanescent fields would also contribute to some of the frustrated light. In practice, the first type of frustration is determined by the surface quality of the optical window. The system can actually benefit from a low quality sample holder with mild scratches. The second type of frustration contributes to a significant amount of light reaching the sample. With the refractive indices of quartz equal to 1.47, sample (water) equal to 1.33, and the coupling angle is greater than 0.67 (39°), the light will refract out of the glass and facilitate illumination (**SF 2.2(R)**). When the coupling angle is less than 0.67, coupled light would continuously undergo TIR at the glass-sample interface. As the third type of frustration, the evanescent field at the glass-sample interface may also facilitate part of the illumination.

---

**Supplemental Figure 2.2**
Diagrams showing the relationship between the acceptance angle and different interfaces
Left: generic case showing maximal angle of $\theta_1$ that leads to TIR
Right: specific case for quartz showing TIR at the glass-air interface and frustrated TIR at the glass-sample interface.

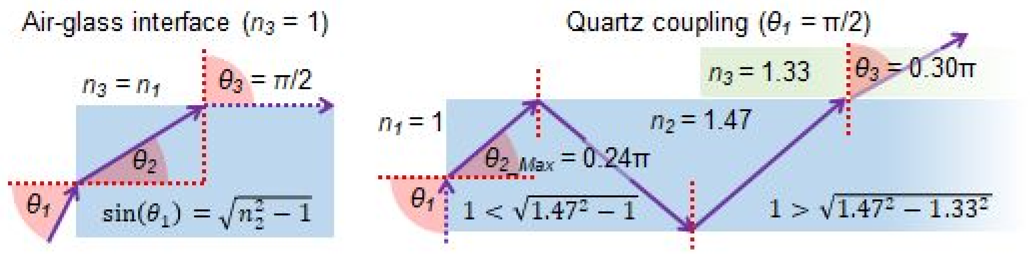

In addition, the relationship between the TIR acceptance angle (y-axis) and refractive indices of the glass substrate (x-axis) is plotted in **SF 2.3** below for an air-glass interface and a sample-glass interface. The TIR acceptance angle, $\theta_{1/TIR}$, is defined by the largest $\theta_1$ that would result in TIR within the glass substrate for corresponding refractive indices across defined interfaces. For parameters above the top (purple, $n_3$=1) curve, TIR would not happen because the coupled light would refract out of the air-glass interface. For parameters below the bottom (blue, $n_3$=1.33) curve, light would undergo TIR at both the air-glass interface and the sample-glass interface. For parameters in between the curves (red area), the light would undergo TIR at the air-glass interface, and refract out at the sample-glass interface. For the special case described in the previous page, when quartz is used as the glass substrate ($n_2$ = 1.47), a large portion of the light would refract out of the glass-sample interface. A small but still significant amount of light would continuously undergo TIR at the sample-glass interface, facilitating illumination through the evanescent field near the glass surface.

---

**Supplemental Figure 2.3**
Plot showing the relationship between the the TIR acceptance angle and different refractive indices for the air-glass interface and the sample-glass interface

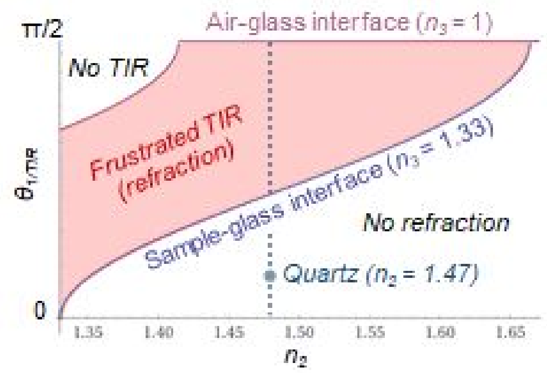

#### 2.2 Total internal reflection (TIR) coupling - interface coupling efficiency

Simple geometric ray tracing discussed in the previous section only partially determines the Pocket MUSE illumination efficiency. In reality, there are two additional major factors affecting the coupling efficiency at this interface in addition to refraction. First, a UVC LED is not a point source. Because UVC LEDs are made on flat sapphire substrate, they usually emit from large flat areas with a nonuniform output profile. Output from a UVC LED is significantly weaker at large output angles compared to small output angles. Second, both reflection and refraction happens at the interfaces of the glass substrate. Considering the Fresnel equation, a significant amount of light undergoes reflection at large output angles instead of refracting into glass substrate. As shown in **SF 2.4**, combining the angular output profile of the LED with the angular coupling efficiency, a significant portion of the light would be coupled into the glass substrate at smaller angles (e.g., <π/4). Coupling in larger angles is not as likely due to low LED output and high reflection.

As discussed earlier in the ray tracing model, the light coupled into the glass substrate within small angles would undergo TIR at both the air-glass and sample-glass interface. Refraction-based frustration would only be facilitated by light entering the glass substrate at the larger angles. Considering the low coupling efficiency of these angles, only a small portion of light would undergo refraction-based frustration. This is not a problem for samples (e.g., whole mount tissues) that are closely attached to the sample holder surface, because they can rely on evanescent field based excitation. However, for samples that are slightly above the sample holder surface (e.g., bacterial suspension), fluorophores outside the evanescent field are only excitable through the refraction-based frustration. Illumination efficiency would be impaired for these types of samples. Still, even though only a small portion of light is coupled in at large angles, low illumination efficiency can be compensated by long exposure times.

---

**Supplemental Figure 2.4**
Combined effects of LED angular output and interface reflection on the overall coupling efficiency

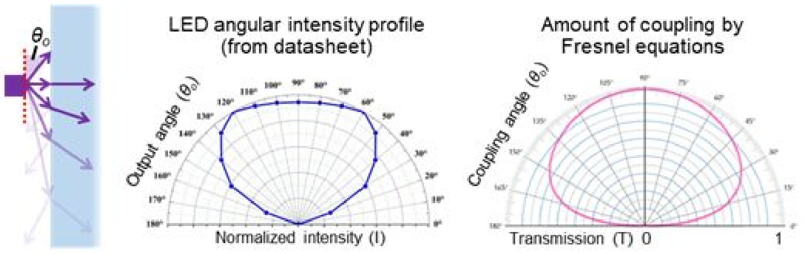


Fresnel equations:

$$T(\theta_o) = 1 - \frac{1}{2}\left( \frac{n_1\cos(\theta_o) - n_2\sqrt{1-\left(\frac{n_1}{n_2}\sin(\theta_o)\right)^2}}{n_1\cos(\theta_o) + n_2\sqrt{1-\left(\frac{n_1}{n_2}\sin(\theta_o)\right)^2}} + \frac{n_1\sqrt{1-\left(\frac{n_1}{n_2}\sin(\theta_o)\right)^2} - n_2\cos(\theta_o)}{n_1\sqrt{1-\left(\frac{n_1}{n_2}\sin(\theta_o)\right)^2} + n_2\cos(\theta_o)} \right)$$

Overall coupling efficiency:

| $\theta_o$ | 15 | 30 | 45 | 60 | 75 | 90 |
|---|---|---|---|---|---|---|
| $I(\theta_o) * T(\theta_o)$ | 0.15 | 0.46 | 0.81 | 0.96 | 0.91 | 0.91 |

#### 2.3 Angled coupling interface

To increase the overall coupling efficiency and refraction-based frustration, an advanced approach is to introduce a smaller angle ($\theta_4$) between the side wall and the top surface of the glass substrate as shown in **SF 2.5** below. This approach could shift the angular distribution of light coupled into the glass substrate, allowing a greater portion of the coupled light to frustrate out of the glass when reaching the sample. With a smaller angle more light escapes at the air-glass interface, but a much higher percentage of light undergoes refraction-based frustration at the sample-glass interface compared to the larger angle. The overall refraction-based illumination efficiency is improved.

By geometric ray tracing, we show that as the angle between the side wall and the top surface of the sample holder decreases ($\theta_4$), $\theta_{1/TIR}$ (the largest $\theta_1$ causing TIR within the glass substrate) also decreases. Assuming the refractive index of the glass substrate is equal to 1.47, the relationship between $\theta_{1/TIR}$ and $\theta_4$ is plotted in the figure below (SF. 2.4) for both the air-glass interface (top blue curve, $n_3 = 1$) and the sample-glass interface (bottom purple curve, $n_3 = 1.33$).

Reducing $\theta_4$ can increase refraction-based frustration at both the glass-air and glass-sample interface. By picking an appropriate reduction of $\theta_4$, one can optimize the amount of light reaching the sample. In the conventional coupling ($\theta_4 = \pi/2$) setup, all the light coupled into the glass substrate will reflect at the air-glass interface and partially refract out at the sample glass-interface. However, because coupling into the glass is inefficient when $\theta_1$ is greater than ~$\pi/4$ (or $\theta_0 < \pi/4$ see SF. 2.3), By reducing $\theta_4$, one can decrease $\theta_{1/TIR}$ and change angular distribution of light entering into the glass substrate. Even though reductions in $\theta_4$ increase the amount of light refracted out at the glass-air and glass-sample interface, $\theta_4$ angles can be chosen to have a greater increase of refraction at the glass-sample interface. We found that a small $\theta_4$ angle (e.g., $\theta_4$ = 1.3-1.5 or 75°- 85°) can get the increased light at the sample with minimal loss at the glass-air interface.

---

**Supplemental Figure 2.5**
Example and ray tracing calculation for angled coupling interface

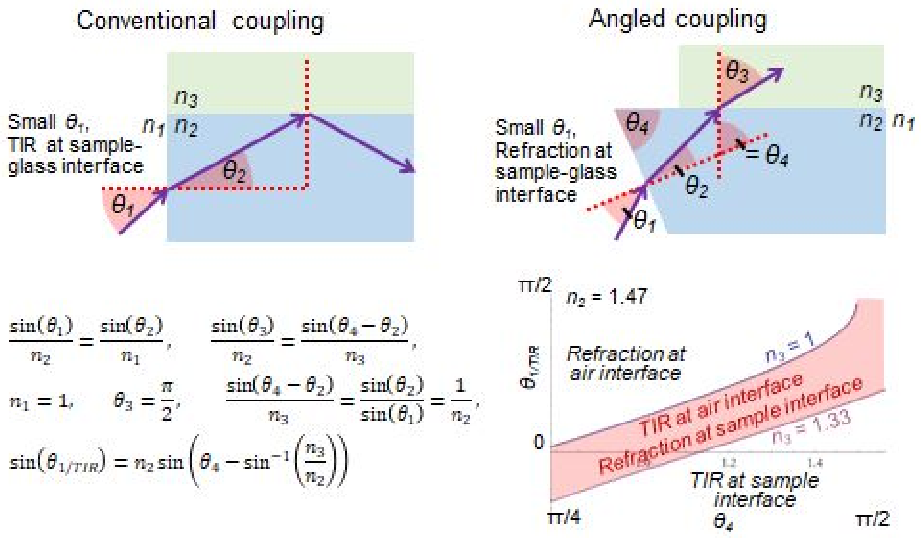

It requires no additional materials and tools to create an angled coupling interface on the side face of the sample holder optical window. However, the fabrication process of the new design is significantly more labor intensive and time consuming compared to the basic design. To precisely polish the optical window at a desired angle, it is necessary to 3D print a polishing holder. As shown in the figure below (**SF 2.6**), the polishing holder is a cuboid with an angled ($\theta_4$) slot, allowing a tight fit of the optical window. It may take a few draft 3D prints to fine tune the width and depth of the slot to ensure proper fit of the optical window. If the slot is too wide, the optical window may slip during the polishing process. If the slot is too narrow, the optical window can easily fracture during the fitting process. In our practice, the dimension tolerance is well below 50 µm and the holder deforms during the polishing process. Also, to prevent excessive scratching during the polishing process, it is necessary to cover the optical window with protection tape. This introduces additional variability to the thickness. Therefore, it is necessary to print and calibrate the polishing holder for each type of optical window. Alternatively, it is possible to print a few polishing holders with slightly different slot widths.

With a fine tuned polishing holder, the polishing process is similar to the process described in the **sample holder** section. The optical window is first gently polished with 300-1500 grit sandpaper, followed by a series of polishing with finer lapping films.The bottom of the polishing holder is set parallel to the sanding paper during the polishing process. It is worth noting that hand polishing is not accurate and can lead to some small curvatures near the edge of the polished surfaces. This variability has minimal negative effect on the coupling performance.

It is necessary to orientate the LED at the same angle in the base plate. Therefore, the LED PCB slots in the base plate should also be tilted as shown in the diagram below (**SF 2.6**). It is slightly more difficult to design and fabricate a base plate with a tilted slot, but the overall prototyping process is similar to the basic base plate when a 3D printer is used. However, introducing two special angled slots may significantly complicate other types of manufacturing processes such as CNC machining and injection molding. Therefore, we only recommend this advanced design consideration for lab scale small volume production.

---

**Supplemental Figure 2.6**
Example of 3D printed polishing holder for creating an angled coupling interface and the corresponding base plate design with tilted LED PCB slots

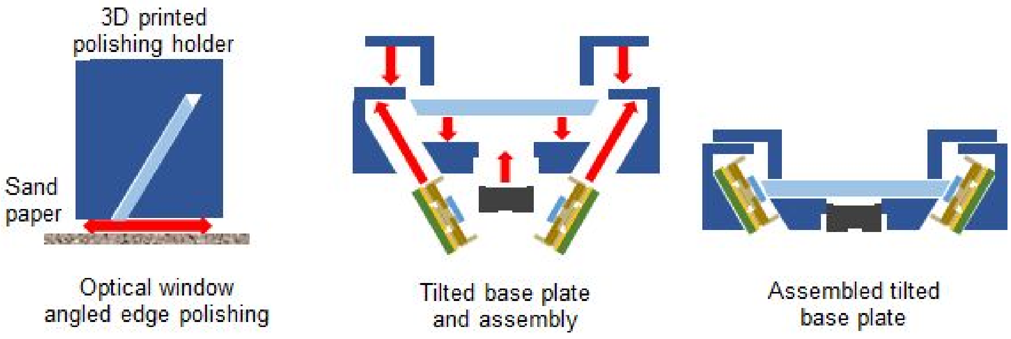

#### 2.4 LED triggering

The LED triggering mechanism is an important design consideration for Pocket MUSE. Continuously driving the LEDs will drain the battery quickly. UVC LEDs in the current Pocket MUSE design do not have sufficient heat resistance for continuous driving. In addition, when the simple constant voltage driving strategy is implemented, UVC LEDs are susceptible to damage due to excess current if kept on for a long period of time. To avoid these problems, the LEDs should only be triggered during image preview and acquisition.

As described in the **electronic** section, the UVC LED is triggered by a mechanical pushbutton switch. When a simple constant voltage supply (without enabling mechanisms) is used, the pushbutton switch connects the LEDs and the power supply. In cases where a more complex driving circuit (e.g., a voltage/current regulator with an enable signal input) is used, the LED triggering switch can be used to enable the power supply directly. This may reduce the risk of power surging that could potentially damage the UVC LEDs.

For advanced designs of Pocket MUSE, there are also several LED triggering channels that do not depend on an external mechanical pushbutton switch. If UV illumination preview is not required, it is possible to use the volume control button on the smartphone to trigger the UVC LEDs. For most smartphones, the volume control buttons can trigger the camera and also produce a trigger signal through the USB output, which can be used to enable the LED driver.

Most camera apps on the smartphone can enable the integrated flashlight during image acquisition. This generates an optical signal that is synchronized with image acquisition, which could be used to trigger the external UVC LED. To implement this strategy, the flashlight of the smartphone needs to be covered by a light shield to prevent stray light leakage into the smartphone camera. A photodiode, photoresistor or phototransistor is installed in the light shield to convert the flightlight signal into an electrical signal, which can be used to trigger the LED driving circuit through the enable signal input.

Alternatively, most smartphones can create a wireless (e.g., Bluetooth or Wifi) flashlight trigger output that can be used to trigger the UVC LEDs in the Pocket MUSE, although this would require more complicated software and hardware development.

#### 2.5 Adjustable focusing

The sanding based alignment procedure is a simple and cost effective way to align the RACL in Pocket MUSE. However, this design has a fixed focal plane. It is necessary to rebuild a new base plate whenever the focus of the device needs to be adjusted (e.g., after a sample holder replacement). Also, because the sanding based alignment requires a significant amount of manual work, it is not suitable for medium to large scale production. In situations where the focus needs to be adjusted regularly, it is necessary to add focus adjustment functionality to Pocket MUSE.

Here, we further investigated and validated two additional focus adjustment strategies. The first strategy is rather simple, but costly to implement. RACLs are made with fine screw mechanisms, which are used to fine align the axial location of the RACLs in smartphone cameras. The screw mechanisms can be directly implemented in Pocket MUSE, avoiding the sanding-based alignment procedure. However, most RACLs have uncommon fine screw pitch sizes (e.g., M6*0.35). Tap tools for these special sized screws are difficult and expensive to obtain. Also, because the optics are highly sensitive to tilt due to the relatively large FOV, it is necessary to ensure the tapping process is as perpendicular as possible to the base plate. Therefore, high precision machining tools and workmanship are required.

The second strategy is to mimic a conventional microscope by introducing a manual linear stage to Pocket MUSE (**SF 2.7**). This design enables precise and real-time focal adjustment during the normal operation of Pocket MUSE. Because the location of the RACL is usually fixed with respect to the smartphone camera, the linear stage could only be implemented to manipulate the position of the sample holder. Therefore, the sample holder is isolated from the base plated in a separate translatable top plate. As shown in the figure supplemental 2.6, the top and base plates are aligned to each other through 4 simple shaft/bearing rails and pulled towards each other through 2 extension springs. The base plate can be axially positioned through a fine adjustment screw. Although this design is functionally more robust, it requires more sophisticated tools and prototyping skills to manufacture. The cost is significantly (>20 times) higher than the basic Pocket MUSE design.

---

**Supplemental Figure 2.7**

The linear stage based Pocket MUSE adjustable focusing mechanism

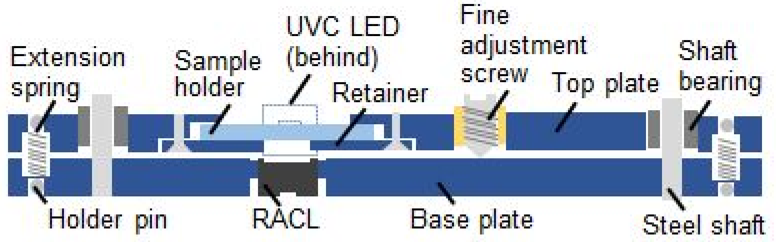


Design concept of the adjustable focusing mechanism

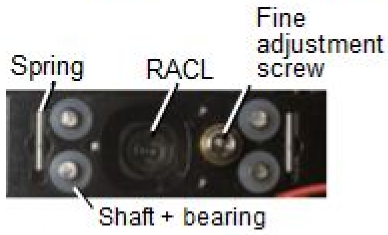


Top view of a adjustable focusing prototype

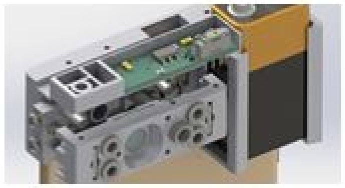

CAD rendering of an adjustable focusing prototype with an LED driver circuit

#### 2.6 Smartphone/camera selection

While Pocket MUSE works with most state-of-the-art smartphones, it is necessary to understand the specifications of the smartphone cameras in order to choose appropriate smartphones for specific microscopy applications.

For common smartphone cameras, the most important specification for Pocket MUSE is the sensor format. As discussed in the result section, improved resolution and magnification of Pocket MUSE depend on a larger image conjugation ratio (e.g., 1:2). Although the ratios are directly determined by the integrated cameras in the smartphones, information about these lenses are usually limited. However, assuming the same field angle, focal length of a smartphone camera lens is usually proportional to its sensor size. Therefore, sensor size becomes the key specification for users to determine the zoom performance of Pocket MUSE. We recommend using smartphone camera sensors larger than 1/3.2”, which provides ≥2X magnification with a 1/7” RACL.

In addition to the sensor size, pixel resolution (sampling) also determines the practical resolution of the system. Assuming the field of view is around 1.5 mm, to achieve theoretical Nyquist sampling for 2 μm optical resolution, about 1500 pixels are required in each dimension. A standard 12 MP (4290 x 2800) sensor is about the appropriate size to meet this requirement. Such a sensor is common in most newer smartphones after 2015 (e.g., iPhone 6). If higher resolution is desired, it is possible to select a smartphone with a larger pixel format. For instance, Samsung developed a 108 MP (12032 x 9024) smartphone camera sensor in 2019, which can provide sampling down to 0.16 μm assuming a 1.5 mm FOV. This may provide additional benefit to compensate for random noise (e.g., hot pixels) and downsampling caused by Bayer filtering.

Another useful specification is the “equivalent focal length”. One way to categorize smartphone cameras is to compare them to standard photographic cameras. Smartphone manufacturers often report “millimeter equivalent” of the smartphone cameras. You may find something like “30 mm equivalent” in the spec sheet of a smartphone, which indicates that the smartphone takes photos similar to a 30 mm compound lens in a standard full frame camera. This information is useful to estimate the magnification of Pocket MUSE with different smartphone cameras. For instance, the 4.25 mm wide angle lens of Apple iPhone Xs is equivalent to a 26 mm compound lens in a standard camera. Comparing this to the 33 mm equivalent camera in an iPhone 6s, the theoretical magnification and resolution of the newer module can be ~20% worse. However, newer smartphones are often equipped with multiple cameras, including telephoto cameras with larger equivalent focal lengths. For example, the telephoto lens of iPhone Xs is 52 mm equivalent, leading to >50% higher theoretical magnification and resolution than 33 mm equivalent cameras. In other words, using a smartphone with a telephoto lens can be a useful way to further improve the magnification and resolution of Pocket MUSE (as shown in **SF 4.7**).

The gap width between a smartphone camera lens and its protection window also determines the performance of Pocket MUSE. With the RACL microscopy setup, it is necessary to keep the entrance pupils of the lenses as close as possible. As the RACL is moved further away from the smartphone camera lens, the effective aperture at large field angles starts to decrease. This can cause significant vignetting and resolution degradation, resulting in reduced FOVs and image quality. This specification is usually not reported or documented for most smartphones, but it could be easily estimated by visual inspection and tends to be smaller in thinner smartphone designs. To achieve the best performance near the edge of the FOV, it is also possible to remove the camera protection window of the smartphone, implementing Pocket MUSE even closer to the smartphone camera.

Finally, it is advantageous to use a smartphone with firmware that supports advanced camera settings and RAW data output. Many old smartphone cameras (e.g., iPhone 5) are designed to mainly serve the default camera apps. Users do not have direct control of many useful camera settings (e.g., analog gain/ISO, shutter speed, etc.). In addition, while many smartphone camera sensors have 12 or 16 bit pixel depth, image data are almost always converted to an 8 bit lossy RGB JPEG format by the default camera apps. Useful information from high bit depth resolution and dynamic range is lost in the process. Recently, many newer smartphones (e.g., iPhone 6 and after) started to offer advanced programming interfaces for 3rd party camera apps. Developers are given more access to the camera configurations, including 12 or 16 bit raw data output. Although the default camera apps remain simple, users can control advanced camera parameters and output lossless data in RAW format. This allows users to set up the smartphone camera to meet the needs of more advanced Pocket MUSE applications.

### 3. Using Pocket MUSE

A brief operation guideline for Pocket MUSE is provided in the material and method section of the main manuscript. However, we were not able to cover some important details due to the word limit. In this section, we aim to provide additional information about the staining strategies and data processing procedures for the general usage of Pocket MUSE. We hope this information can inspire Pocket MUSE users to develop customized staining protocols and data processing pipelines to meet the need of specific imaging tasks.

#### 3.1 Staining and dye selection

The extremely simple staining procedure is a major highlight for many Pocket MUSE applications. Here, we provide additional insight about how a staining solution can be formulated for Pocket MUSE applications. In general, dyes are dissolved in solvent containing water and alcohol (e.g., methanol, ethanol or isopropanol). For most common histology dyes, 30-70% alcohol is a good starting point. It helps to both solubilize the dye molecules and permeabilize the tissue. Depending on the chemical properties of the dyes, tap water (e.g., polarized dyes), denatured alcohol (e.g., lipophilic dyes) or 1-10% DMSO solution can be used as alternatives. When necessary, pH of the solvents may be adjusted for specific dyes using concentrated buffer solutions at the desired pH. The same solvents could also be used to wash the dyes from the stained samples, although tap water washing is sufficient for most types of samples we have tested. For dyes that only fluoresce strongly when binding to certain structures (e.g., DAPI), the washing step could be skipped.

Because Pocket MUSE simultaneously acquires fluorescence images in 3 spectral channels, channel crosstalk can be significant compared to conventional fluorescence microscopy. If two dyes have spectral emission closer to each other, it can be difficult to unmix them from the adjacent color channels. For the best image contrast, 2 dyes each emitting in the blue and red channels are recommended (e.g., DAPI + rhodamine). When formulating a staining solution, it is also necessary to take the intrinsic fluorescence and autofluorescence of the sample into consideration. For instance, many fixed animal tissue samples have a relatively strong paraffin induced fluorescence in the blue-green channels. Therefore, dyes with strong emission and high structural affinities, such as DAPI and Alexa 488 conjugated antibodies, are critical for generating good contrast in these channels.

In addition to the fluorescent properties, the absorption/chromatic properties of the dyes also need to be considered, because absorption also affects the spectral property of the dyes. In fact, many applications can benefit from this combined fluorescent and chromatic contrast. For instance, similar to the hybrid mode imaging, it is possible to use a fluorescent dye to create a global background emission, and then use an absorptive dye to generate chromatic contrast on the background. One example is starch grains stained with iodine water, which has high absorption throughout the spectrum and appear black in the image even under UVC excitation. It is also possible to use an absorptive dye to block out-of-focus light and improve image contrast. Adding 0.1% Sudan Black B or Light Green SF in a staining solution can effectively block light for a typical DAPI/rhodamine stain (**Fig. 2e**).

The necessary duration for staining is usually several seconds for most samples and staining solutions we have tested. Most fluorescent dyes only require a dip in the staining solution to effectively stain the surface of the sample. Some chromatic dyes may require longer staining time (e.g, iodine). When a continuous supply of water is available, excess staining solution on the sample can be directly washed out with flowing water for ~10 s. When the amount of washing solvent is limited (e.g., when tap water is not available), we recommend first absorbing excess staining solution with a piece of absorbent material, then soaking the sample in the washing solution for ~30 s.

All solvents mentioned above have relatively low toxicity at the working volume. However, dye staining is usually difficult to remove from skin and clothes. Therefore, gloves are recommended when handling these solutions. High concentration alcohols are highly volatile, so it is necessary to keep the staining solution in sealed containers. Some solvents are flammable. It is important to prevent fire hazard during the staining process, but the risk of fire is low at the working volume.

The fluorescent emission of most common histology dyes are not reported below 300 nm. Therefore, we tested the performance of a range of common histology dyes under a 285 nm UVC LED using an RGB camera. For reference, basic information and UVC excited fluorescence of these dyes are shown in **SF 3.1** below:

---

**Supplemental Figure 3.1**
UVC fluorescence, chemical/biochemical properties and molecular structures of common dyes. Each dye was blotted on a piece of cotton (not fluorescent) and exposed to 285 nm light and imaged with a RGB camera. For each dye, an image is shown. Next to the image, the name, concentration, exposure time, average RGB values, physical properties, and chemical structure are provided.

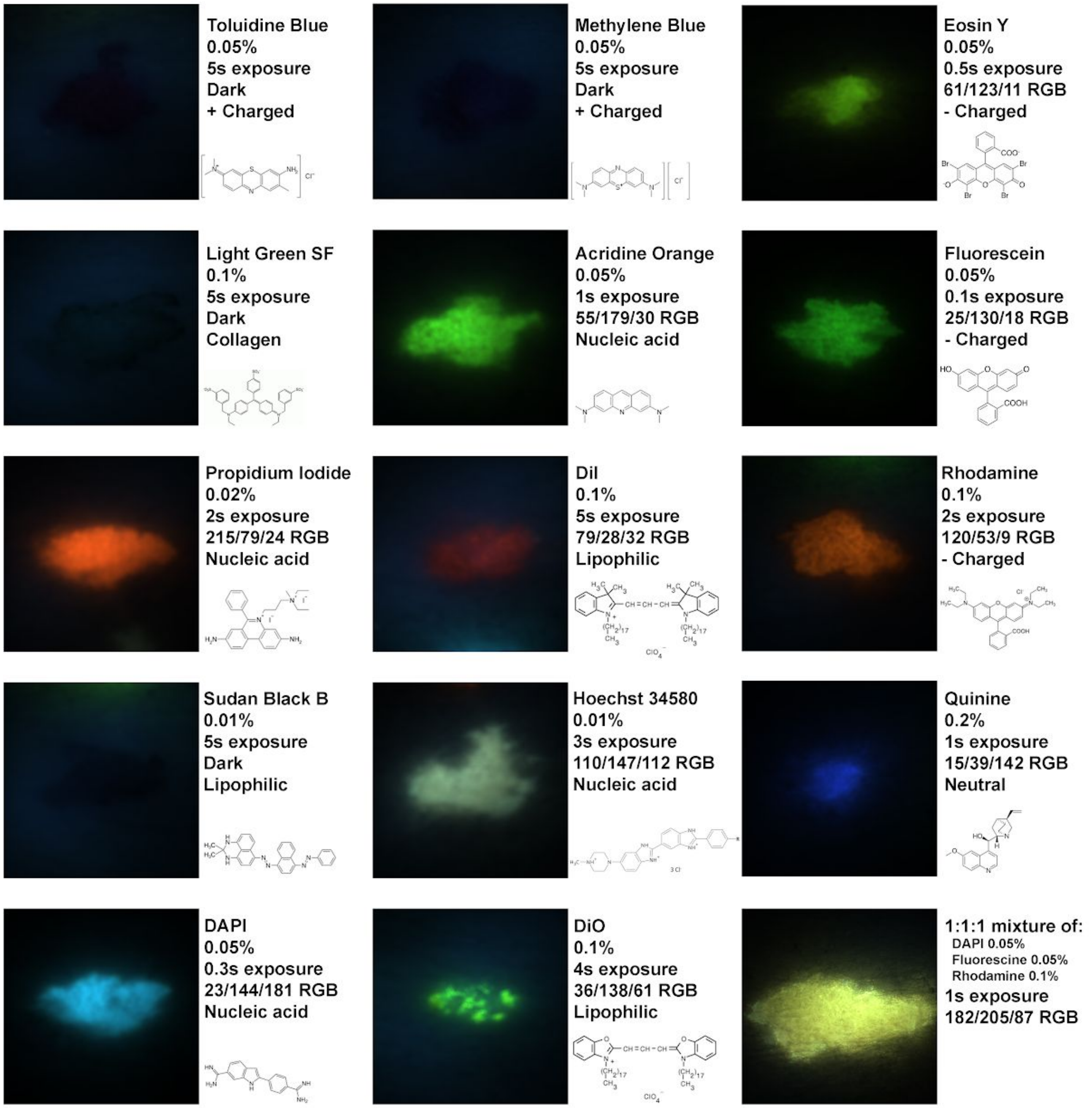

#### 3.2 Sample loading and unloading

After staining and washing, excess water on the sample surface needs to be removed before the sample is loaded onto the Pocket MUSE sample holder. However, it is also important to avoid drying the sample surface completely, because a small amount of liquid provides the necessary surface tension to keep the sample attached to the sample holder. Drying samples too much can also introduce a significant amount of air bubbles between the sample and the sample holder, degrading the image quality. Instead of using a piece of dry absorbent material (e.g., a cotton pad), it is easier to first wet the material with the washing solvent.

In our current Pocket MUSE system, it is recommended to limit one of the sample dimensions to 3 mm. With a 0.5 mm thick sample holder (optical window) and 2 LEDs on opposite sides, samples wider than 3 mm can cause significant nonuniform illumination. Defining $\theta_2$ as the angle at which a ray is coupled into the glass, and t as the thickness of the glass (see **SF 3.2** left), the ray can only reach $2t*\cot(\theta_2)$ from the edge of the sample assuming it only reaches the glass-sample interface once. We define $\theta_{2\_Max}$ as the maximum angle that a ray could enter the glass substrate, and $\theta_{2/CRIT\_water}$ as the minimal $\theta_2$ that could result in TIR at the glass-sample interface (see **SF 3.2** right). Assuming all rays enter the glass at random angles and locations, a relatively uniform illumination could be achieved within $2t*\cot(\theta_{2\_Max})$ from the edge of the sample. Beyond $2t*\cot(\theta_{2\_Max})$, the intensity of the light starts to decay. And beyond $2t*\cot(\theta_{2/CRIT\_water})$, no ray could reach the sample as the glass-water critical angle is reached.

The ray tracing calculation only represents the ideal condition where the light is uniformly distributed throughout the angles. In reality, light is never uniformly distributed across the angles. To understand this effect in practical terms, we conducted additional optical simulation and experiments. As shown in **SF 4.8** and **SF 4.9**, decaying of the illumination from the edge of the sample is observed in both simulation and experimental setups. This nonuniform illumination is compensated using two LEDs from opposite sides of the sample.

Samples can be loaded and unloaded from the Pocket MUSE sample holder using tweezers. The sample should be removed from the device before completely drying, especially when a volatile solvent is used. It can be difficult to remove completely dried samples from the sample holder, causing damage to the optical window. To prevent scratching the surface of the sample holder, plastic tweezers are recommended. If the sample holder surface needs to be cleaned after unloading the sample, 70% ethanol or isopropanol on a Q-tip can effectively remove residue debris and dyes, and disinfect the sample holder surface.

**Supplemental Figure 3.2**
Ray tracing schematic showing the decay of illumination from the edge of the sample.

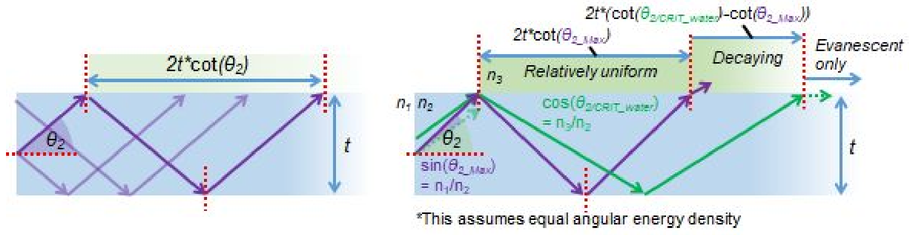

#### 3.3 Data acquisition

There are a few different ways to acquire Pocket MUSE data after the sample is loaded on the sample holder. The simplest way is to use the default camera app of the Android or iOS system. While some Android phones provide a “manual mode” in the default camera apps, the iOS camera app only offers limited access to manual camera parameter adjustment. If camera parameters are adjustable, the user can preset the exposure time, focus, ISO (gain) and white balance to desired values for Pocket MUSE imaging. The UVC LEDs can be switched on occasionally to help adjust these numbers. Good image contrast can be achieved with a manual setup of the camera. If the camera parameters are not adjustable, the user needs to keep the UVC LED on in the preview mode and let the camera automatically adjust these image parameters. Because Pocket MUSE data often have high contrast in at least one color channel, auto focusing and brightness can still create acceptable images for general use. Still, image quality can be compromised because the algorithms are not designed for microscopy imaging. It may also drain the battery much faster because of the extended use of the UVC LEDs.

It is helpful to use 3rd party camera apps to access advanced parameters of the smartphone cameras when they are not adjustable by the default app (e.g., iOS). These camera apps are particularly useful for saving the data in camera RAW format, a function that most default camera apps do not have. For example, the interface of a 3rd party camera app (Halide - available on both iOS and Android) is shown in the figure below, which provides a convenient way to adjust camera parameters that are important to Pocket MUSE imaging. For advanced users, it is also possible to program customized camera apps with advanced data processing capabilities.

Camera RAW data often have sensor readout linearly mapped to pixel values. When background autofluorescence contributes to a great portion of the emitted signal, the image contrast can be affected in the RAW data. For better data visualization directly on the smartphone, it is necessary to process the data to enhance the image contrast. This could be done directly on the smartphone by adjusting the brightness/contrast (linear stretch), gamma (exponential fitting), shadow/highlight (partial histogram stretch) and hue/saturation (color rebalance) using the default photo editing apps. These adjustments would generate a non-linear mapping between the raw data values and displayed pixel values, improving the image contrast.

**Supplemental Figure 3.3**
Left: Screenshot of a 3rd party camera app (Halide), showing a brain slice imaged with Pocket MUSE
Right: A comparison between an unprocessed image as shown on the smartphone screen and a contrast enhanced image showing clearer contrast of the neural fibers in the green channel.

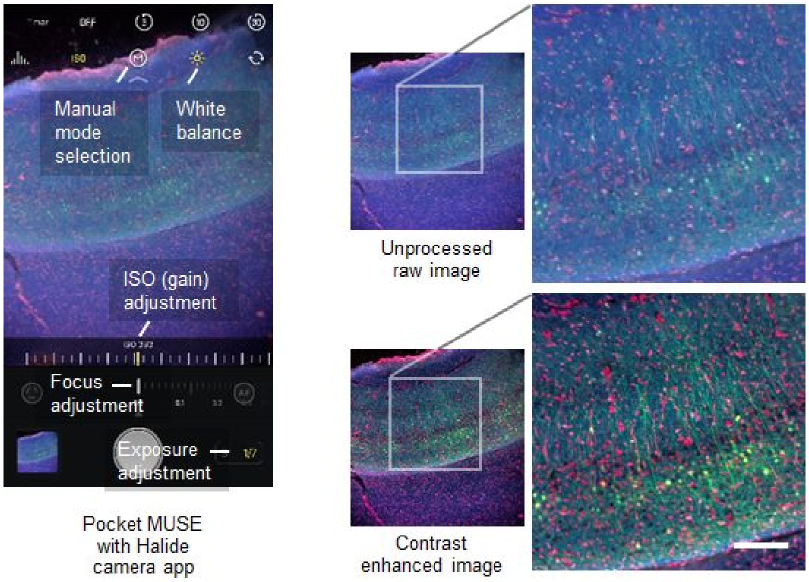

##### 3.4 Raw data conversion

The camera RAW data are the sensor readouts saved as single channel arrays. RGB channels are not separated. Unprocessed RAW data appear as grayscale images, and the original Bayer RGB pattern is clearly shown when the images are zoomed in (as shown in **SF 3.4** below). This type of data can be readily processed with image processing algorithms after the channels are resampled. The pixel values are proportional to the photon energy emitted from each pixel location.

To properly visualize the images, they first need to be converted into a standard RGB format. In addition, as the full dynamic range of the image data is well preserved in the RAW files, most of the time only a small portion of the range contains useful information. Depending on what dyes are used, it is necessary to enhance the contrast of corresponding channels in order to correct the white balance of the image. For instance, with propidium iodide and Alexa Fluor 488 staining, most fluorescent signals are emitted at the green and red channels. Auto contrast enhancement (linear histogram stretch) by linearly remapping 99% of the pixels between value 1 and 255 often provides an acceptable contrast for data visualization.

However, because there are 2 green channels in each 2x2 Bayer unit matrix, and there are spectral overlaps between adjacent color channels. Also, for different image sensors, Bayer filters can have varied spectral distributions. Thus, the simple resampling based data conversion discussed above is not the most efficient way to convert RAW camera data into RGB format, causing degraded resolution and contrast. For more efficient RAW data conversion, it is recommended to use a professional raw data converter such as Adobe CameraRaw or RawTherapee, especially when the data is used for direct visualization. These tools can perform semiautomated raw data conversion by applying preset camera profiles/settings using advanced algorithms. In addition, users without extensive image processing experience can adjust the image contrast parameters through interactive toggle bars and real-time preview.

---

**Supplemental Figure 3.4**
Example images showing unprocessed RAW data (top) of a brain slice stained and a zoomed-in image clearly showing the Bayer RGB pattern. The RAW data is converted to RGB format without further processing as shown in the bottom left, and linearly contrast enhanced as shown in the bottom right.

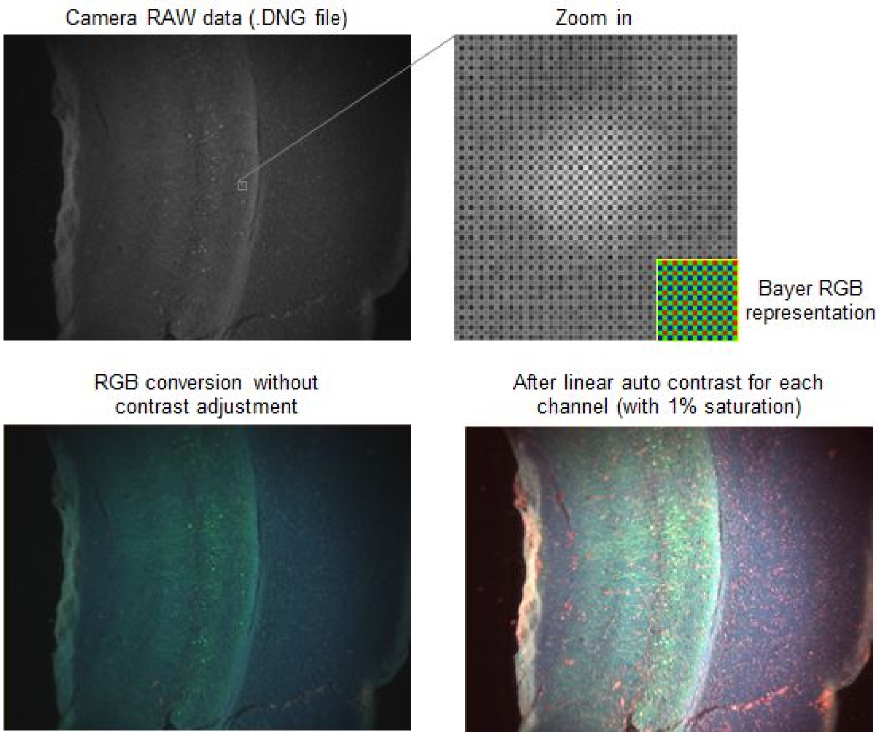

For images acquired for quantitative assessment, it is possible to process the RAW files directly with scientific image processing tools such as ImageJ/FIJI[8] or programming languages such as Python and Matlab. Direct opening of camera raw data is often not supported by the default programs. It is necessary to use 3rd party data loading software such as the OME Bio-Formats [9] plugin/library. An example ImageJ Macro code for converting camera RAW files into RGB format is shown below:

---

**Example Code 1**

```
// ImageJ Macro code for isolating RGB channels from raw data
imgId = getImageID();
isolateCh("red", 0, 0);
selectImage(imgId);
isolateCh("greenA", 0, -1);
selectImage(imgId);
isolateCh("greenB", -1, 0);
selectImage(imgId);
isolateCh("blue", -1, -1);

// Combine isolated channels and convert to RGB
run("Merge Channels...", "c1=red c2=greenA c3=blue create");
run("RGB Color");

// Isolate channels based on their pixel coordinates
function isolateCh(title, x, y) {
  imgW = getWidth() / 2;
  imgH = getHeight() / 2;
  run("Duplicate...", "title= &title");
  run("Translate...", "x=&x y=&y interpolation=None");
  run("Size...", "width=&imgW height=&imgH interpolation=None");
}
```

For enhanced data visualization, most images demonstrated in this manuscript are converted from camera RAW data using a professional RAW data conversion tool called Adobe Camera Raw. An example interface of this program is shown in **SF 3.5** below. There are many image contrast parameters that can be controlled. In brief, temperature and tint adjust the color (white) balance of the RGB channels. Exposure and contrast each controls the offset and slope of the linear mapping between the raw pixel value and displayed pixel value. Highlights, shadows, whites and and blacks each applies a nonlinear mapping to the histogram, enhancing the contrast of a small portion of the displaced dynamic range. Texture enhances the high frequency features in the image. Clarity and dehaze reduces low frequency background light in the image. Vibrance and saturation controls the hue of the image. Although optimal values of these parameters vary for images acquired under different setups, it is important to keep all the settings constant when comparing images for scientific studies.

**Supplemental Figure 3.5**

Example screenshots showing the Adobe Camera Raw 11.4 interface.
Top: A RAW data file loaded into the software with the camera default profile setting;
Bottom: The same file with enhanced contrast after some parameter adjustment.

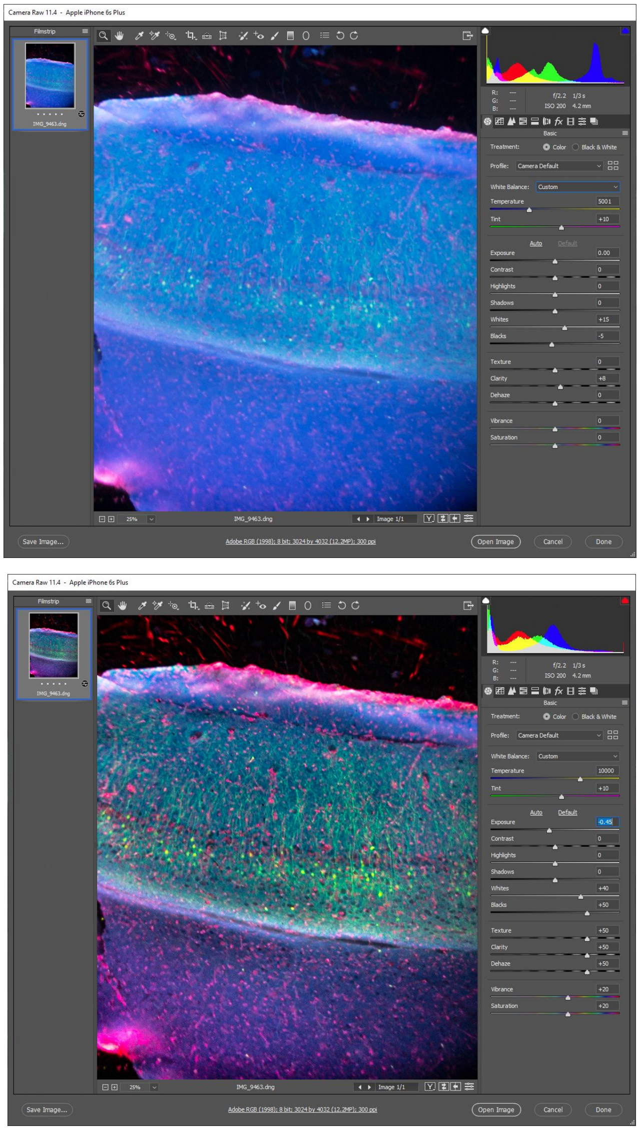

Adobe Camera Raw is a commercial software that can be costly to some users. An alternative RAW conversion software is RawTherapee (**SF 3.6**). This software is free and open source, providing similar functionality to Adobe Camera Raw.

**Supplemental Figure 3.6**
Example screenshots showing the RawTherapee 5.8 interface.

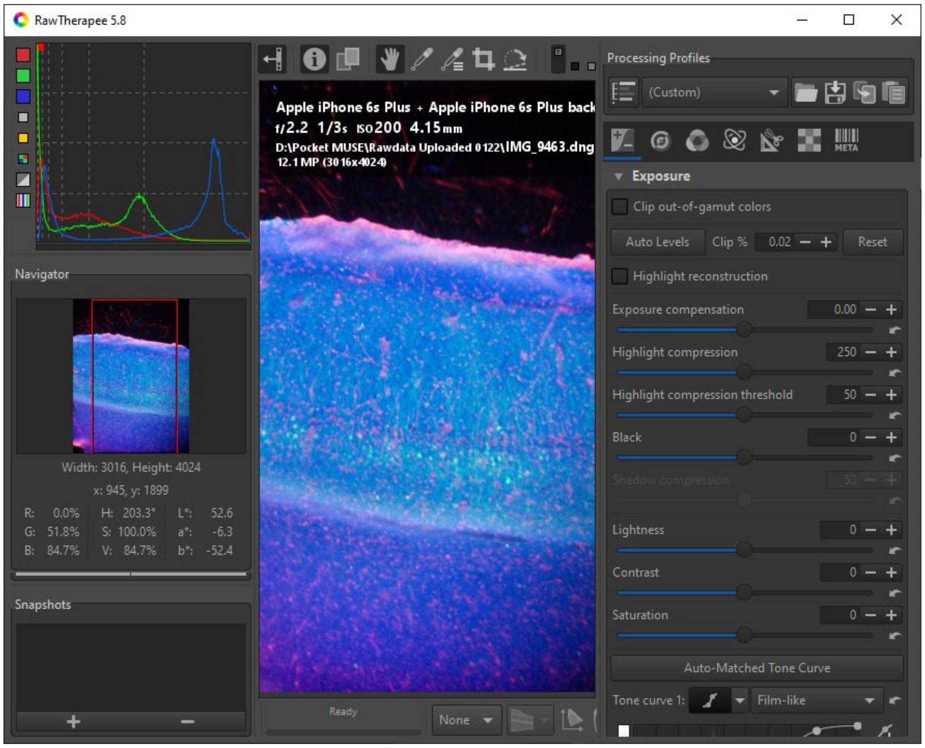

#### 4. Additional characterization and images

**Supplemental Figure 4.1**
Resolution measurement at different locations of the FOV.
Setup: iPhone 6s+ (4032 × 3024 RGB sensor) + 1/7” RACL
Relatively uniform resolution (USAF 1951 Group 9-1, <1 μm) could be achieved across the entire FOV.

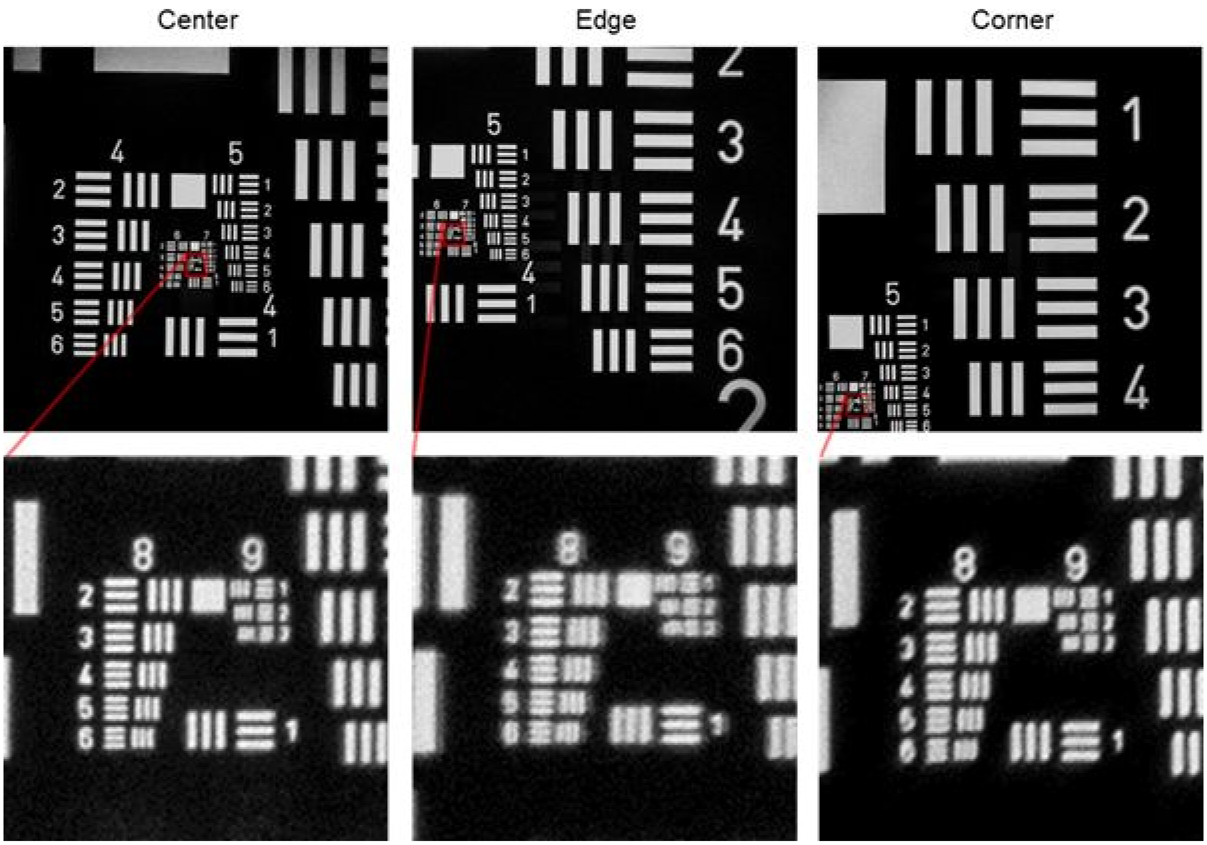

**Supplemental Figure 4.2**
Resolution comparison between Pocket MUSE and a benchtop microscope.
Left: iPhone 6s+ (4032 × 3024 RGB sensor) + 1/7” RACL
Middle: Keyence BZ-X810 (1920 x 1440 monochrom camera) + Nikon Plan Apo 10X/0.45 DIC N1 objective
Right: Keyence BZ-X810 (1920 x 1440 monochrom  camera) + Nikon Plan Fluor ELWD 20X/0.45 objective

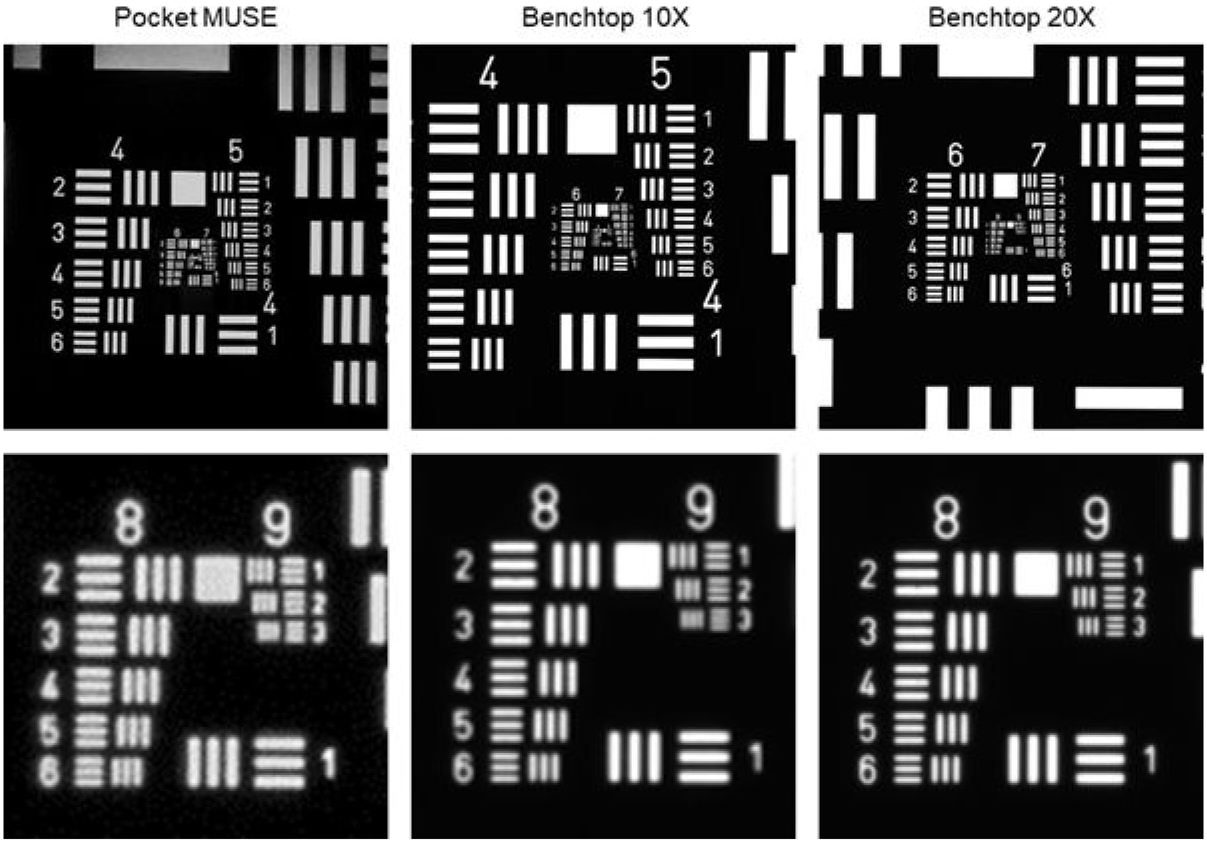

**Supplemental Figure 4.3**
Resolution measurement w/ iPhone 6s+ and 4 different RACL
(Left: full FOV / Right: center 10% FOV)

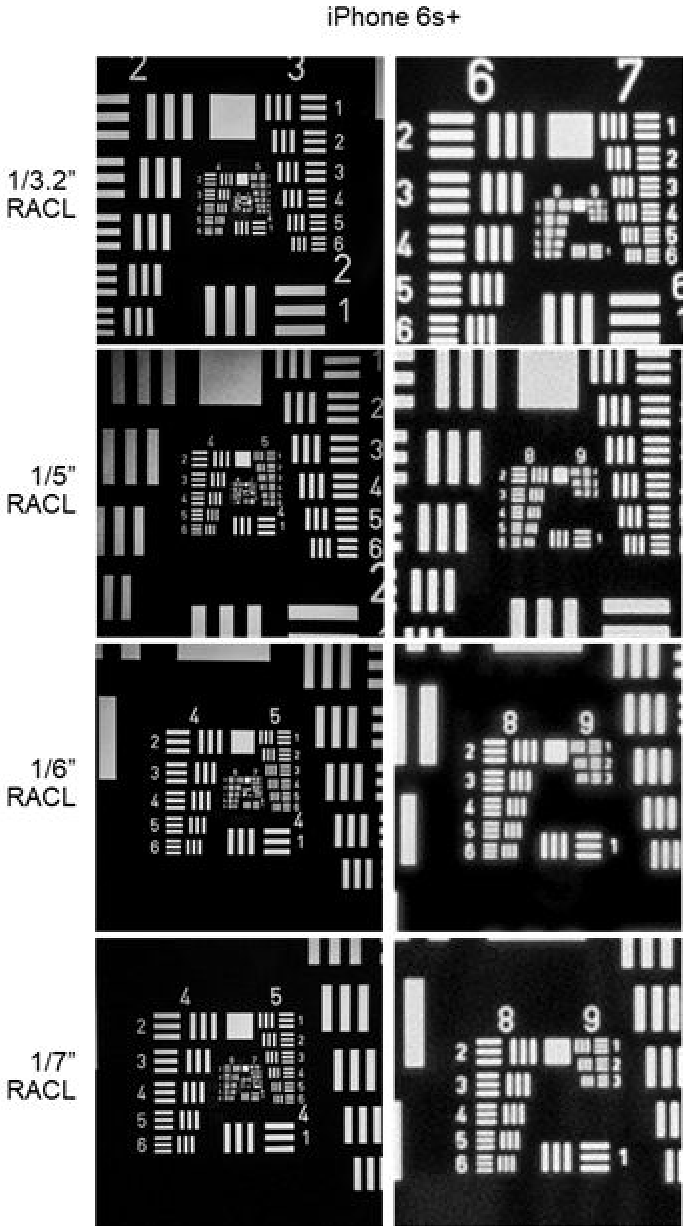

**Supplemental Figure 4.4**
Resolution measurement w/ iPhone X (1X camera) and 4 different RACL
(Left: full FOV / Right: center 10% FOV)

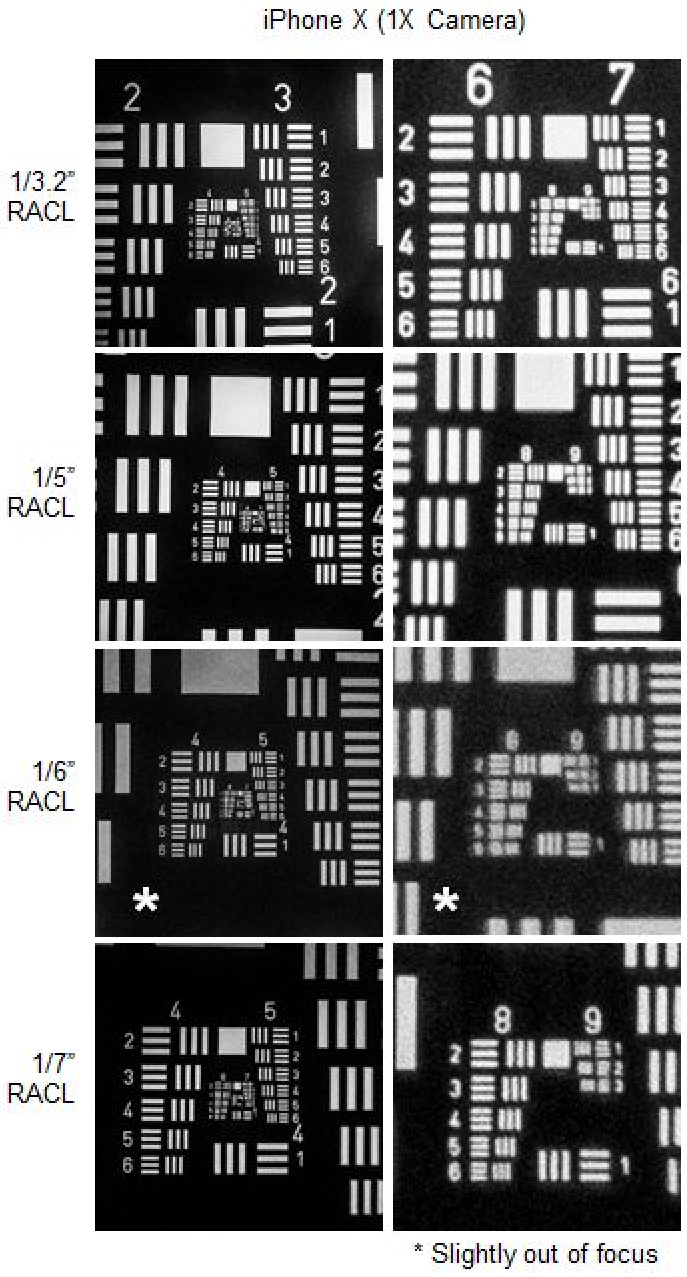

**Supplemental Figure 4.5**

Resolution measurement w/ iPhone X (2X camera) and 4 different RACL
(Left: full FOV / Right: center 10% FOV)

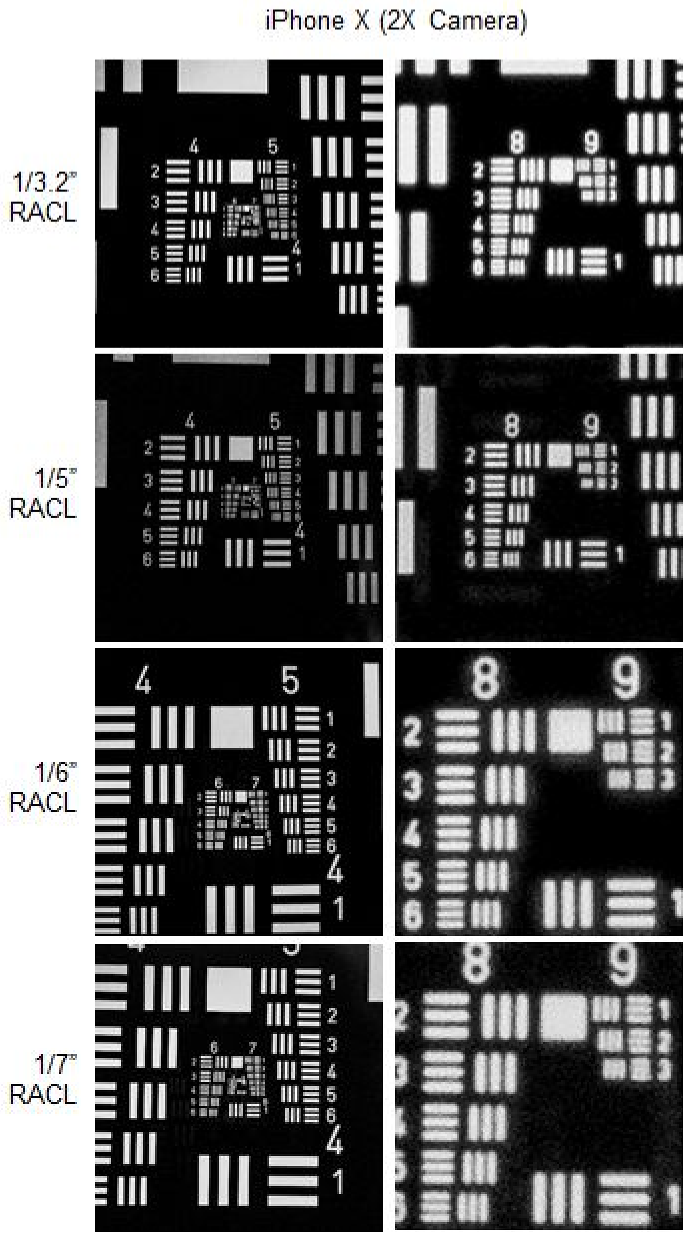

**Supplemental Figure 4.6**
Resolution measurement w/ MI6 (1X camera) and 4 different RACL
(Left: full FOV / Right: center 10% FOV)

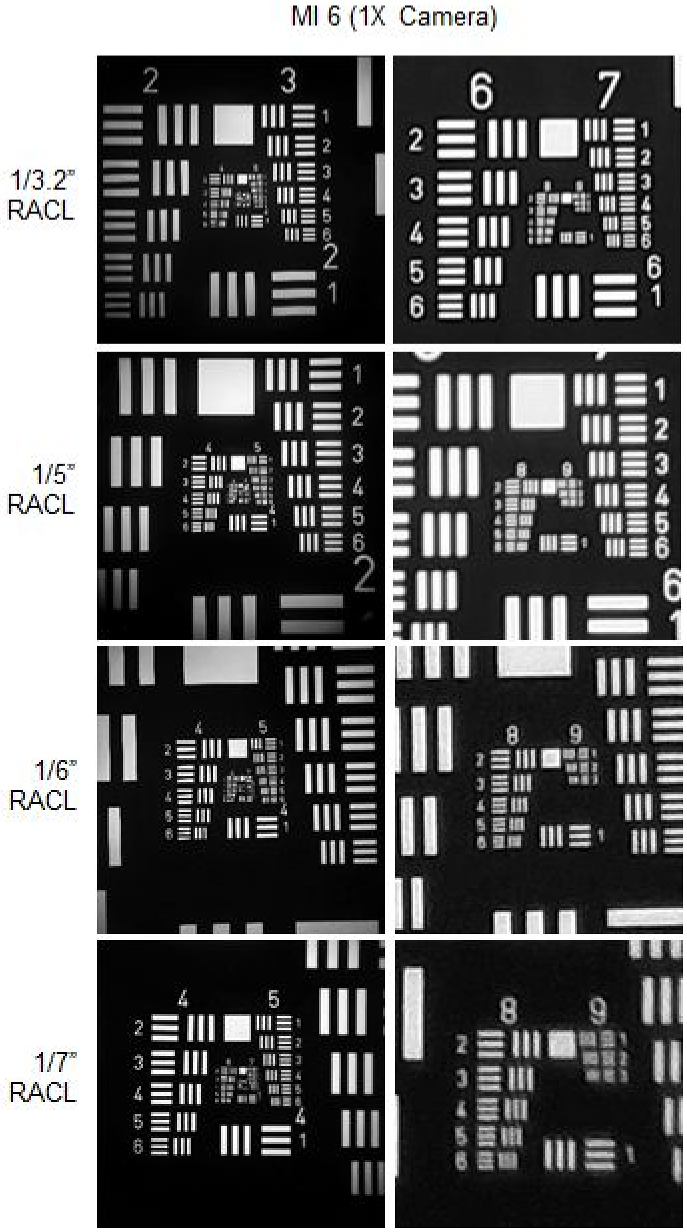

**Supplemental Figure 4.7**
Resolution measurement w/ MI6 (2X camera) and 4 different RACL
(Left: full FOV / Right: center 10% FOV)

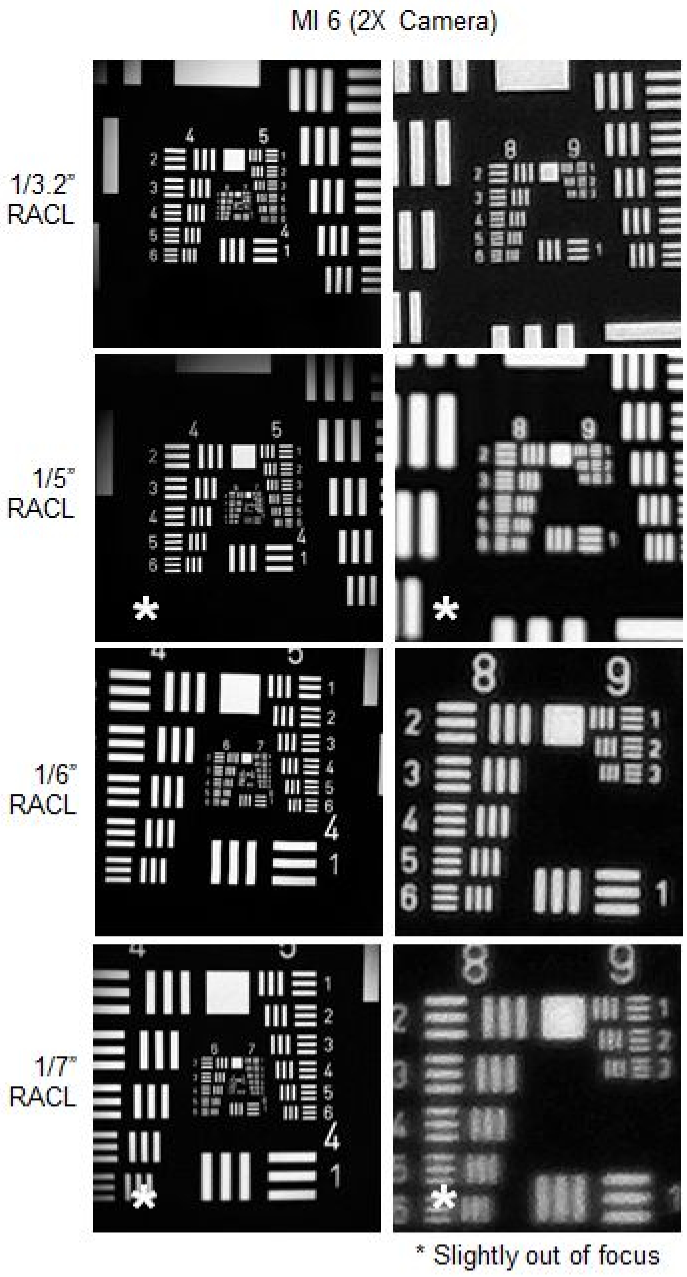

**Supplemental Figure 4.8 Difference between the single-LED and dual-LED setups**
Zemax non sequential ray tracing showing the difference between using one LED and using two LEDs.
a. Schematic of ray tracing. The gray blocks represent the sample holder, the orange squares represent the sample, and the purple lines represent the rays coming out of the LEDs;
b. Irradiance maps across the sample holder top surface, all light is absorbed by the sample in the center and does not show up in the irradiance map;
c. Irradiance map at the sample showing a significant decay from right to left with a single LED and a relatively uniform distribution of light with two LEDs at opposite sides;
d. 1D plots of the irradiance across the dotted black lines in c.

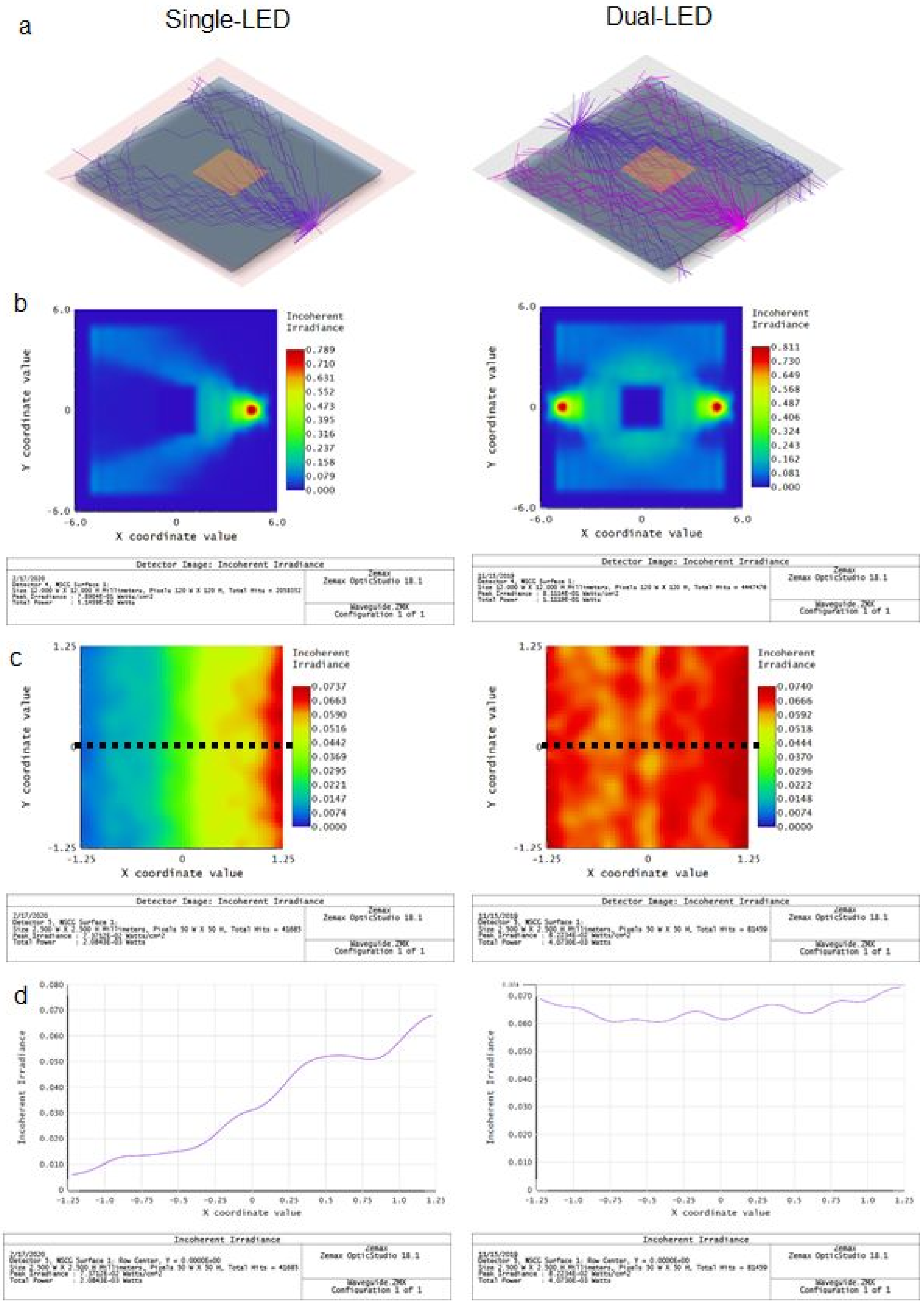

**Supplemental Figure 4.9 Difference between various sample sizes**

Zemax simulation and experimental data showing the illumination uniformity with 3 different sample sizes

a. Irradiance maps across the sample holder top surface, all light is absorbed by the sample in the center and does not show up in the irradiance map;

b. Irradiance map at the sample, showing slightly higher (for 2x2 mm), relatively uniform (for 2.5x2.5 mm), and lower (for 3x3 mm) irradiance at the center of the sample;

c. 1D plots of the irradiance across the dotted black lines in b;

d. Experimental images showing the irradiance of different sized sample phantoms (fluorescent sticky note) on a real Pocket MUSE device showing relatively uniform illumination for 2x2 mm and 2.5x2.5 mm phantoms, and dimmed illumination at the center of the 3x3 mm phantom, confirming the simulation results.

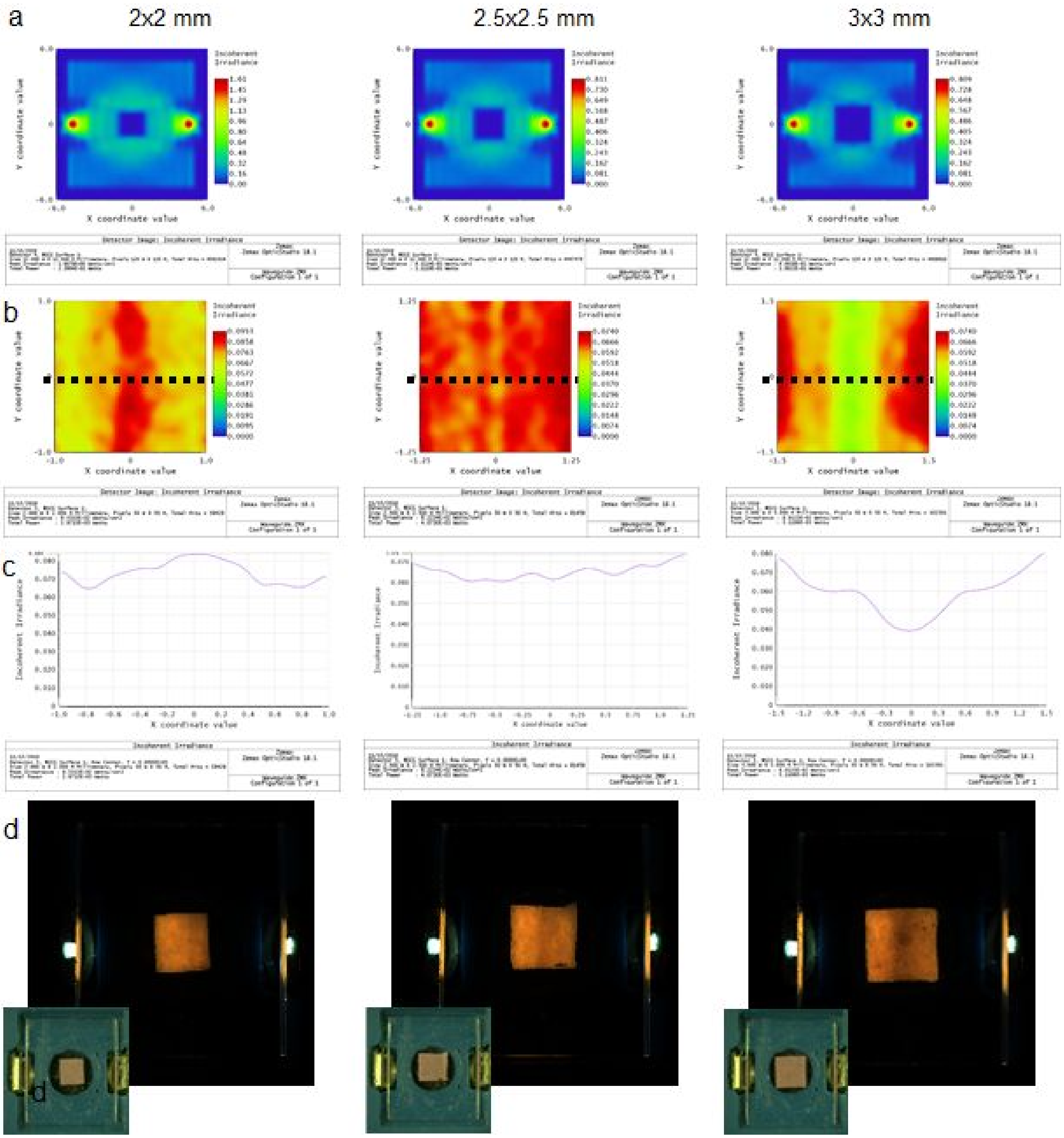

**Supplemental Figure 4.10**
Additional images showing the difference between the bright-field (top) and the MUSE illumination (bottom). scale bars: 300 μm

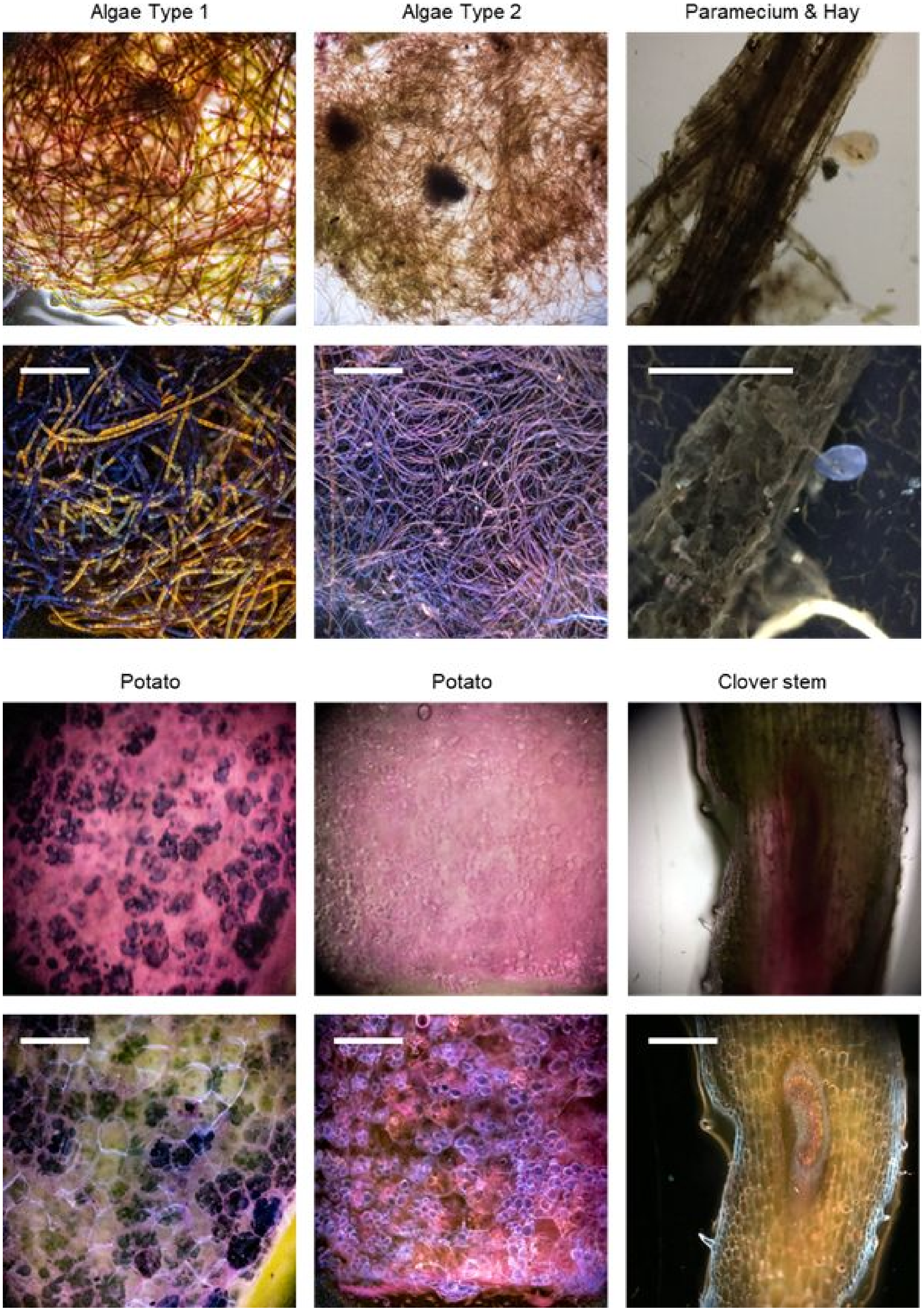
